# Developmental stage-dependent decoupling of molecular and phenotypic responses to heat in wheat grain

**DOI:** 10.64898/2026.07.16.733488

**Authors:** Farhad Masoomi-Aladizgeh, Thomas M. Ashhurst, Lake-Ee Quek, Yasmin Asar, Chad L. Moore, Samiuela Lee, Anowarul I. Bokshi, Errin Johnson, Samson Dowland, Michelle L. Wood, Kevin Begcy, Ben Crossett, Ali Khoddami, Richard Trethowan, Daniel K.Y. Tan, Brian J. Atwell, Thomas H. Roberts

## Abstract

Heatwaves during flowering and grain development threaten global wheat production, yet the extent to which developmental stage shapes molecular and phenotypic responses remains unclear. Here, we investigated whether heat stress (36/29°C for 48 h) imposed at closely spaced developmental stages surrounding anthesis generates stage-specific molecular responses that are associated with subsequent effects on grain properties. Heat exposure increased floret abortion most strongly when it was imposed at the trinucleate stage (TN; ∼48%) compared with the binucleate (BN) and early post-anthesis stages. Methylation levels were largely unaffected by prior heat treatments; however, heat induced stage- and locus-specific methylation changes, particularly at BN. Spatial RNA-seq revealed tissue-specific heat responses that were distinctively different in BN and TN. Integrated metabolomic analyses revealed stage-dependent metabolic reprogramming, including shifts towards stress-associated pathways. Despite heat stress at BN generating broader molecular reprogramming in developing grain, exposure at TN produced stronger effects on grain set and composition, revealing a decoupling between the magnitude of responses and phenotypic outcomes. Together, these findings demonstrate that subtle differences in developmental stage influence the complex molecular responses to heat and subsequent grain properties.

## Introduction

Heatwaves during flowering increasingly threaten global wheat (*Triticum aestivum*) production, with current varieties showing limited resilience to moderate temperature increases (Xiong and Reynolds 2024). Across major wheat-growing regions, each 1℃ rise in mean growing-season temperature is estimated to reduce yields by ∼6% (Asseng et al. 2014). Wheat is particularly vulnerable during its reproductive phase, when even brief heat exposure can cause severe damage to developing florets. Temperatures of 35℃ applied around meiosis can induce complete sterility (Draeger and Moore 2017), while prolonged heat at anthesis can lead to yield reductions of up to 38% (Mirosavljevic et al. 2024). Grain quality—including traits such as protein composition—is also susceptible to high temperatures. Most controlled-environment studies have focused on the grain-filling period as the key window during which elevated temperatures alter final grain composition (Hafeez et al. 2023), yet much earlier reproductive stages may also play a critical role, but the impact has remained underexplored.

As heatwaves become longer, hotter, more frequent, and earlier in the growing season (Jyoteeshkumar reddy et al. 2021), they increasingly coincide with these highly susceptible stages, underscoring the need for mechanistic insight to support breeding for reproductive heat tolerance. Importantly, development along the wheat inflorescence is asynchronous—between spikelets, among florets, and even within anthers—complicating precise assignment of heat impacts on grain traits (Masoomi-Aladizgeh et al. 2024). This complexity impedes accurate staging, isolation, and molecular profiling of anatomically defined pollen precursors and developing grains, ultimately slowing gene discovery for heat tolerance.

Across several crops, male gametophytes demonstrate pronounced thermal susceptibility, with heat disrupting pollen development and dramatically reducing yields (Begcy et al. 2019; Hu et al. 2024; Masoomi-Aladizgeh et al. 2020). In wheat, however, it remains unclear whether transient heat stress at specific reproductive stages triggers distinct, stage-dependent molecular responses that ultimately influence grain yield and quality. Elucidating whether heat responses differ across these tightly timed stages is therefore essential for resolving the temporal specificity of reproductive heat sensitivity and its downstream effects on grain development and quality.

Despite extensive studies on heat stress during early reproductive development, it remains unclear how developmental stage influences the propagation of stress-induced molecular changes in subsequent grain development, and how these responses are spatially organised within grain tissues. We hypothesised that (i) heat stress imposed at closely spaced developmental stages surrounding anthesis generates distinct molecular signatures during grain development, and (ii) the magnitude of molecular reprogramming predicts downstream phenotypic outcomes. To test these hypotheses, we integrated spatial transcriptomics, epigenetic profiling and metabolomics across reproductive stages surrounding anthesis to determine how developmental stage shapes the propagation of heat-induced molecular responses into grain development.

## Materials and methods

### Plant material and growth conditions

Wheat plants (*Triticum aestivum* L., 2*n* = 6*x* = 42) were grown in controlled environment rooms at the University of Sydney. A heat-tolerant line known for strong heat tolerance and grain quality, PBI13505 (KAUZ/PASTOR//PBW343/3/KIRITATI/4/FRNCLN/5/SUNTOP), developed by the University’s Plant Breeding Institute (Fig. S1), was studied. Seeds were sown in 17 × 18 cm pots containing 1.5 kg of premium plus superior potting mix (Scotts, Osmocote^®^) supplemented with ∼20 g of slow-release fertiliser, Osmocote (NPK 11:1:3). Plants were maintained at 22/15 ± 2°C (day/night), 12/12 h light/dark, and a minimum of 400 µmol m^−2^ s^−1^ photosynthetically active radiation. Fertiliser was applied weekly as 0.8 g L^-1^ Aquasol^®^ (NPK 23:3.95:14; Yates, Australia), and plants were drip-irrigated to ensure adequate water supply. Each pot contained three seedlings and was treated as one biological replicate. The experimental design is illustrated in Figure 1.

**Figure 1.**
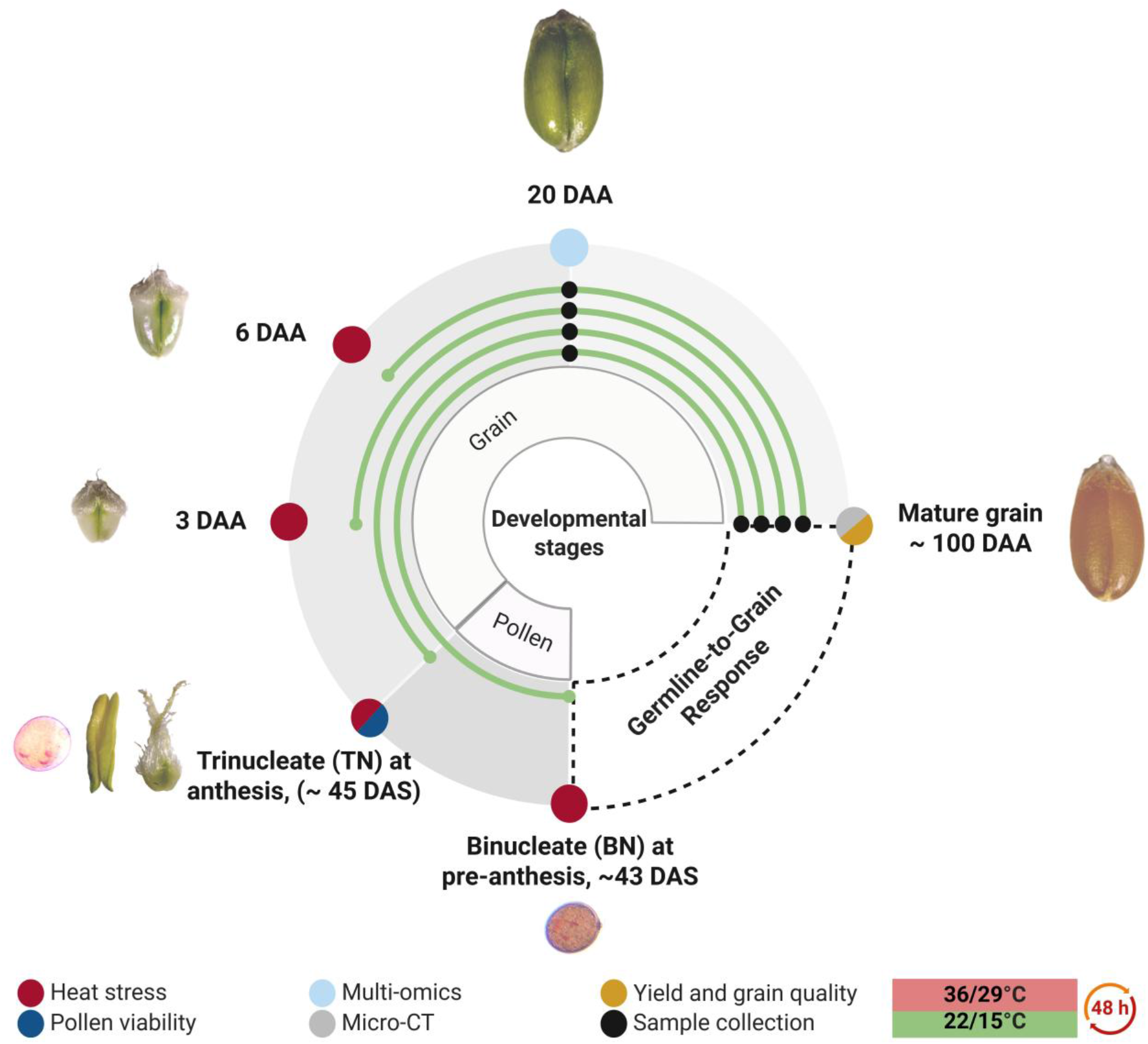
Experimental design, illustrating how increased temperatures during pollen development, anthesis, and post-anthesis were interrogated for their effects on immature and mature wheat grains. Stages during pre-anthesis (binucleates, approximately 43 days after sowing, DAS), anthesis (trinucleate, approximately 45 DAS), and post-anthesis (3 and 6 days after anthesis, DAA) were subjected to heat stress (36/29°C for 48 h). Subsequently, samples were collected at 20 days after anthesis (DAA) and at maturity for various analyses.

### Elevated temperature treatments

Plants grown under controlled-environment conditions were exposed to heat stress of 36/29 ± 2°C (day/night) for 48 h at either the binucleate (BN) stage, anthesis (trinucleate, TN), or early grain-filling stages, including 3 and 6 days after anthesis (DAA). This brief heat treatment encompassed late pollen development, fertilisation, and early endosperm formation. Plants were returned to ambient conditions after heat exposure. Primary tillers that initiated anthesis before completion of heat treatment were excluded. From each primary tiller, up to eight central spikelets were collected from both control and heat-treated plants for molecular analyses. Only the primary and secondary florets were harvested; all tertiary and quaternary florets were discarded. To assess long-term impacts of heat applied at different developmental stages, immature grains at 20 DAA were sampled from plants heated at the binucleate stage (IM-BN), anthesis (IM-TN), 3 DAA (IM-3D), and 6 DAA (IM-6D). For grain-yield and quality analyses, mature grains from the same heat-treatment windows (M-BN, M-TN, M-3D, M-6D) were collected from entire spikelets across the inflorescence, including grains from the third and fourth florets.

### Determining pollen and grain developmental stages

The morphological development of the wheat plants was monitored regularly to accurately identify the stages of pollen and grain development for heat stress treatments. To target a defined stage of pollen development, heat was applied when approximately half of the inflorescence had emerged from the flag-leaf sheath (∼2 days before anthesis), ensuring that the primary and secondary florets in the middle region of the spike had reached the binucleate stage. This approach accounted for the inherent asynchrony of wheat inflorescence development—and the resulting variation in heat-stress susceptibility—as described by Masoomi-Aladizgeh et al. (2024). To confirm developmental stages, anthers were squashed on a glass slide to release pollen, which was then stained with 2% acetocarmine to visualise nuclei. These observations were used to infer staging in the remaining plants without dissection, thereby avoiding disruption to inflorescence growth. To define grain-filling stages, florets undergoing anthesis were marked individually, and grain development was monitored from anthesis onward to accommodate floret-level asynchrony (Masoomi-Aladizgeh et al. 2024).

### Pollen viability analysis

Pollen viability was assessed using impedance flow cytometry (IFC) on an Ampha^TM^ Z32 instrument (Amphasys AG, Switzerland), following Bokshi et al. (2021). In brief, two yellow, non-dehisced anthers from individual florets were placed into 1 mL of AmphaFluid 6 buffer (Amphasys AG, Switzerland) in a microcentrifuge tube. The anthers were crushed to release pollen, and the suspension was filtered through a 70-µM mesh into a 5-mL sample tube. The final volume was adjusted to 2 mL before loading onto a 120-µM microchip (Amphasys AG, Switzerland). Measurements were performed at 12 MHz for 2 min and analysed using AmphaSoft (v 2.1.6.0). Pollen viability was reported as a percentage.

### Yield

Mature grains from control and heat-stressed plants were harvested approximately 100 days after sowing. Because M-BN and especially M-TN spikes lacked most primary and secondary florets after exposure to high temperatures, grains from the entire spike were collected for mature-grain analysis regardless of spikelet position or floret number. Grains were further dried at 28°C for 10 h in a forced-air incubator (TLM–570, Thermoline Scientific) before downstream analyses. Yield traits, including grain number per spike, grain weight per spike, and individual grain weight were then recorded.

### Near Infrared (NIR) analysis

NIR scans were conducted using a Bruker MPA FT-NIR spectrometer (Bruker Optik GmbH, Ettlingen) to characterise global compositional variation in grain chemistry. Mature grains were finely milled and sieved using a 200-mesh sieve (74 μm pore size) to remove coarse particles for quality analysis. A total of 100 mg of fine flour was transferred to 8 mm vials (Bruker Optik GmbH, Part No. IN430) and placed on the sphere microsample accessory for analysis. Spectra were recorded from 12,500 to 3,750 cm^-1^ at 4 cm^-1^ resolution, averaging 64 scans per sample, using OPUS (v 8.7.41, Bruker Optik GmbH, Ettlingen). Technical replicates were averaged in OPUS and subsequently analysed using Quasar (v 1.11.1).

### Amino acid analysis

A total of 100 mg of sieved flour (74 μm mesh) was used for amino acid profiling. Samples were hydrolysed in 6 M hydrochloric acid for 24 h at 110℃, during which asparagine and glutamine were converted to aspartic and glutamic acids, respectively. After hydrolysis, an internal standard mix (norvaline and α-aminobutyric acid; Nva/AABA) was added. The hydrolysate was diluted with ultrapure water, and 10 μL was derivatised using an AccQ-Tag Ultra Derivatisation Kit (Waters Corporation, Milford) according to the manufacturer’s instructions. Derivatised samples were analysed on an ACQUITY™ Ultra Performance LC system (Waters Corporation, Milford) following Cohen (2001), with some modifications from Boogers et al. (2008). Separation was performed on an ACQUITY UPLC BEH C18 1.7 µm column (Waters Corporation, Milford) with UV detection at 260 nm. Amino acids were identified and quantified using Empower (v 3.6.1, Waters) against internal-standard calibration curves (0–50 pmol; 1 µL injection). Amino acid composition was expressed as the percentage of each amino acid relative to the total detected amino acids. Samples were analysed as single measurements with three biological replicates.

### X-ray micro-computed tomography (micro-CT)

Micro-CT was conducted to assess the effects of heat stress on grain internal architecture, with additional 3D reconstruction providing context for surface features. Mature grains were immersed in 1 mL of fixative containing 2.5% (v/v) glutaraldehyde + 2% (w/v) paraformaldehyde in 0.1 M phosphate buffer (pH 6.8) for 48 h at 4°C. The fixative was discarded, and samples were rinsed in 0.1 M phosphate buffer (three times, 5 min each), then stored in 0.1 M phosphate buffer at 4°C. Samples were subsequently incubated in 2% osmium tetroxide in Milli-Q water for four weeks, followed by three washes with Milli-Q water.

Samples were mounted in a hydrated state in Eppendorf tubes containing Milli-Q water and imaged using a Bruker SKYSCAN 2214 micro-CT scanner. Projections were acquired at an isotropic voxel size of 2.5 μm using a source voltage of 50 kV and a source current of 165 µA, with no source filter applied. Three-dimensional reconstructions were generated using NRecon software (v 2.3.1.1), and 2D cross-sections and 3D volumes were visualised in DataViewer (v 1.6.0.0) and CTvox (v 3.3.1), respectively. Crease depth (CD) was measured at the distance from the modified aleurone to the midpoint of the distal nucellar projection (NP). Crease width (CW) was measured at 50% of CD along the crease axis, providing a relative measure of internal crease geometry that scales with overall crease elongation.

### Reduced representation bisulfite sequencing (RRBS)

DNA methylation analysis was performed on immature grains (20 DAA) subjected to heat (36/29°C for 48 h) at the binucleate (IM-BN) and trinucleate (IM-TN) stages. Immature grains were preserved in RNAlater^TM^ (Thermo Fischer Scientific, Waltham) at 4°C for up to one week during sampling, after which the RNAlater was discarded and the grains were stored at −80°C. Samples were freeze-dried and then homogenised as described above. Approximately 20 mg of the ground tissue was used for DNA isolation following Masoomi-Aladizgeh et al. (2023).

RRBS library preparation was conducted according to established protocols (Gao et al. 2014; Pan et al. 2018) by CD Genomics (US). Briefly, 1 μg of genomic DNA was digested with MspI (New England Biolabs), end-repaired, A-tailed, and ligated to 5-methylcytosine-modified adapters. Size selection was then performed to isolate 150–350 bp fragments. Bisulfite treatment was performed using the ZYMO EZ DNA Methylation-Gold Kit. The bisulfite-converted DNA fragments were amplified for 12 PCR cycles using 25 μL of KAPA HiFi HotStart Uracil+ ReadyMix (2X) and 8-bp index primers (1 μM). Libraries were quality-checked using an Agilent 2100 Bioanalyzer, quantified with a Qubit dsDNA HS Assay, and sequenced on an Illumina HiSeq platform using paired-end 150 bp reads.

### Spatial transcriptomics

Spatial RNA-seq profiling was performed on immature grains (20 DAA) subjected to heat at the binucleate (IM-BN) and trinucleate (IM-TN) stages. Cross-sections were prepared as described by Masoomi-Aladizgeh et al. (2025). Briefly, immature grains were snap-frozen in liquid-nitrogen-cooled isopentane and stored at −80°C until sectioning. Frozen tissues were incubated in 200 µL of 30% sucrose in 1X PBS containing RNase inhibitor (1 U/µL) for 2 h at 4°C, rinsed with nuclease-free water, briefly dried on Kimwipes, and returned to −80°C. To prepare transverse sections (15 μm), tissues were sectioned on a cryostat (Leica CM3050) at −21°C (specimen temperature −15 to −20°C, depending on sectioning quality) and embedded in optimal cutting temperature (OCT) compound (Tissue-Tek). Sections were stored at −80°C until further processing.

Spatial gene-expression profiling followed the Visium HD 3’ fresh-frozen protocol (10x Genomics) with minor modifications. Slides were incubated at 37°C for 2 min, fixed in 100% methanol (−20°C, 30 min), incubated in ice-chilled 100% isopropanol (1 min, room temperature), and air-dried for 5 min. Throughout tissue preparation, 0.1X PBS was used to prepare solutions at a maximum temperature of 30°C. Libraries were quality-checked using a TapeStation, quantified with a Qubit dsDNA HS Assay, and sequenced at the Ramaciotti Centre in Genomics (Australia) on a NovaSeq X Plus (10B 300-cycle format; 43-10-10-75 bp), with a 2% PhiX spike-in and a loaded library concentration of 150 pM. FASTQ files were generated for downstream analysis.

Sequencing reads were processed using Space Ranger (v 4.0.1; 10x Genomics). A custom wheat reference genome was constructed from IWGSC RefSeq v2.1 obtained from URGI-INRAE (Zhu et al. 2021) using *mkref*, and reads were aligned using *count*, generating spatially resolved UMI count matrices at 8 µm bin resolution. Filtered UMI count matrices were imported into *Seurat* (v 5.3.1) in R using *BPCells* (0.3.1) for memory-efficient processing. Low-quality bins were excluded based on total UMI counts, retaining bins between the 2^nd^ percentile (196 UMIs) and an upper threshold of 500 UMIs prior to downstream analysis. Data were log-normalised, scaled, and subjected to principal component analysis (PCA). Biological replicates were integrated using *Harmony* (v 1.2.4) to correct for batch effects and generate a unified spatial transcriptomic atlas across conditions. Unsupervised clustering was performed using a shared nearest neighbour graph and the Louvain algorithm, with results visualised using Uniform Manifold Approximation and Projection (UMAP). Spatial annotation was guided by marker genes identified using *Seurat*’s FindMarkers, selected based on average log₂ fold change (avg_log2FC ≥ 0.5) and statistical significance (adjusted *p*-value ≤ 0.05). For differential expression analysis, raw UMI counts were aggregated either across all spatial bins within each replicate (sample-level pseudobulk) or within each cluster–replicate combination (cluster-specific pseudobulk). Differential expression analysis (|log₂FC| ≥ 0.5, *p* ≤ 0.05) was performed using *edgeR* (4.0.16) with trimmed mean of M-values (TMM) normalisation and quasi-likelihood testing. Spatial transcriptomics data were visualised in Loupe Browser (v 9.0.0; 10x Genomics).

### Metabolomics analysis

Metabolite extraction was performed following Zhang et al. (2021), with minor modifications. Freeze-dried immature grains (30 h; Alpha 1–4 LSCbasic, CHRIST) were homogenised using a Precellys Evolution Touch homogeniser (10 cycles of 30 s at 6,500 rpm; Bertin Technologies) with five Zirconox beads (2.8–3.3 mm). For extraction, 30 mg of ground tissue was mixed with 800 µL of pre-chilled methanol:water (1:1 v/v), containing internal standards (0.25 µM camphorsulfonate for negative mode; 0.5 µM phenylalanine-D8 for positive mode). Chloroform (800 µL) was added, samples were vortexed for 15 min at 4℃, and then centrifuged at 20,000 *g* for 5 min. The upper aqueous phase was collected and lyophilised in a SpeedVac (Savant SPD140DDA) without heat. Polar extracts were re-suspended in 200 µL of mobile phase (1:1 buffer A:buffer B; buffer A = 95:5 H_2_O:ACN with 20 mM ammonium acetate and 20 mM ammonium hydroxide; buffer B = 100% ACN), centrifuged at 20,000 *g* for 10 min at 4℃, and 100 µL of supernatant was transferred to LC-MS vials.

LC–MS analysis was performed on a Shimadzu LC-40D X3 coupled to a ZenoTOF 7600 (SCIEX). A 5-µL injection was run at 0.25 mL min⁻¹ with the column oven at 30°C and autosampler at 5°C, using a 22-min gradient. The ZenoTOF 7600 operated in both positive and negative modes, using standard vendor settings for source gases and collision energies. Data-dependent TOF-MS/MS spectra were acquired over appropriate *m/z* ranges at standard resolution. Data were acquired in Analyst and processed in MS-DIAL (v5.5.250530). Features were curated (manual reintegration as needed), normalised to the mode-specific internal standards, log_2_-transformed, annotated via MS-FINDER with cross-checks in MassBank/HMDB, and analysed statistically in MetaboAnalyst and R (v 4.5.1) (R Core Team 2023).

### DESI-MS imaging

For desorption electrospray ionisation (DESI) mass spectrometry imaging (MSI), immature grains were snap-frozen in liquid-nitrogen-cooled isopentane and stored at −80°C. To prepare 30-μm cross-sections, Milli-Q water was used instead of OCT as an embedding medium to avoid ion suppression in DESI, then tissues were sectioned on a cryostat (Epredia NX50) at −15°C and thaw-mounted onto plain glass slides. DESI buffer was 98% MeOH: 2% H_2_O: 0.1% formic acid containing leucine enkephalin (80 ng mL⁻¹) as lock mass. Imaging was performed on a DESI-XS source coupled to a SYNAPT G2 (Waters) in positive ion mode over *m/z* 50–1200. The pixel size was 15 µm, and the stage speed was 15 µm s⁻¹. Lockmass calibration was performed using 50 ng mL^-1^ of leucine enkephalin ([M+H]^+^, *m/z* 556.2615). Data were acquired in MassLynx using the DESI source configuration, then processed and visualised in Waters High Definition Imaging (HDI) Software (Waters). Ion images were normalised to TIC prior to mapping the spatial distribution of the ions.

### Statistical analysis

Data were analysed and visualised using R (v 4.5.1) (R Core Team 2023) and GraphPad Prism (v 11.0.0). Normality and homogeneity of variances were assessed using the Shapiro-Wilk and Brown-Forsythe tests, respectively. Depending on data distribution, either a one-way ANOVA or a Kruskal-Wallis test was applied. Post hoc comparisons used Dunnett’s test for parametric data and Dunn’s test for non-parametric data. Differences with *p ≤* 0.05 were considered statistically significant.

## Results

### Developmental checks and sampling considerations

To minimise developmental asynchrony between tillers and among spikelets along the inflorescence, grain samples were collected only from the main tiller and from the primary and secondary florets of approximately six to eight of the central spikelets to avoid top and bottom spikelets that typically develop more slowly (Masoomi-Aladizgeh et al. 2024). Heat treatments (36/29°C for 48 h) were applied at the binucleate (BN), trinucleate (TN), 3 days after anthesis (3 DAA), and 6 days after anthesis (6 DAA) using spikes reliably staged through their emergence from the flag-leaf sheath and microscopy observations. In wheat, TN corresponds to the time that anthesis occurred. Heat treatment at BN started when spikes were approximately half-emerged (∼2 days before anthesis), while TN and grain-filling treatments were initiated using individually labelled florets at anthesis. Spikes that progressed to anthesis during the 48 h BN treatments were excluded to ensure consistent developmental staging.

Heat treatments produced distinct stage-dependent morphological effects on the grain (Fig. 2a–b). Heat at BN and TN resulted in visibly larger immature grains at 20 DAA, and this size difference persisted to maturity. Heat stress at these reproductive stages also led to increased floret abortion, particularly at TN. By contrast, heat at 3 or 6 DAA did not induce abortion; spikelets generally retained up to three developing florets, comparable to control plants. However, grains appeared greener (more advanced) than controls at corresponding time points, suggesting accelerated maturation under the heat stress. For mature-grain analyses, entire spikes were harvested across all treatments to ensure sufficient grain, particularly at BN and TN, for which abortion reduced grain number.

**Figure 2.**
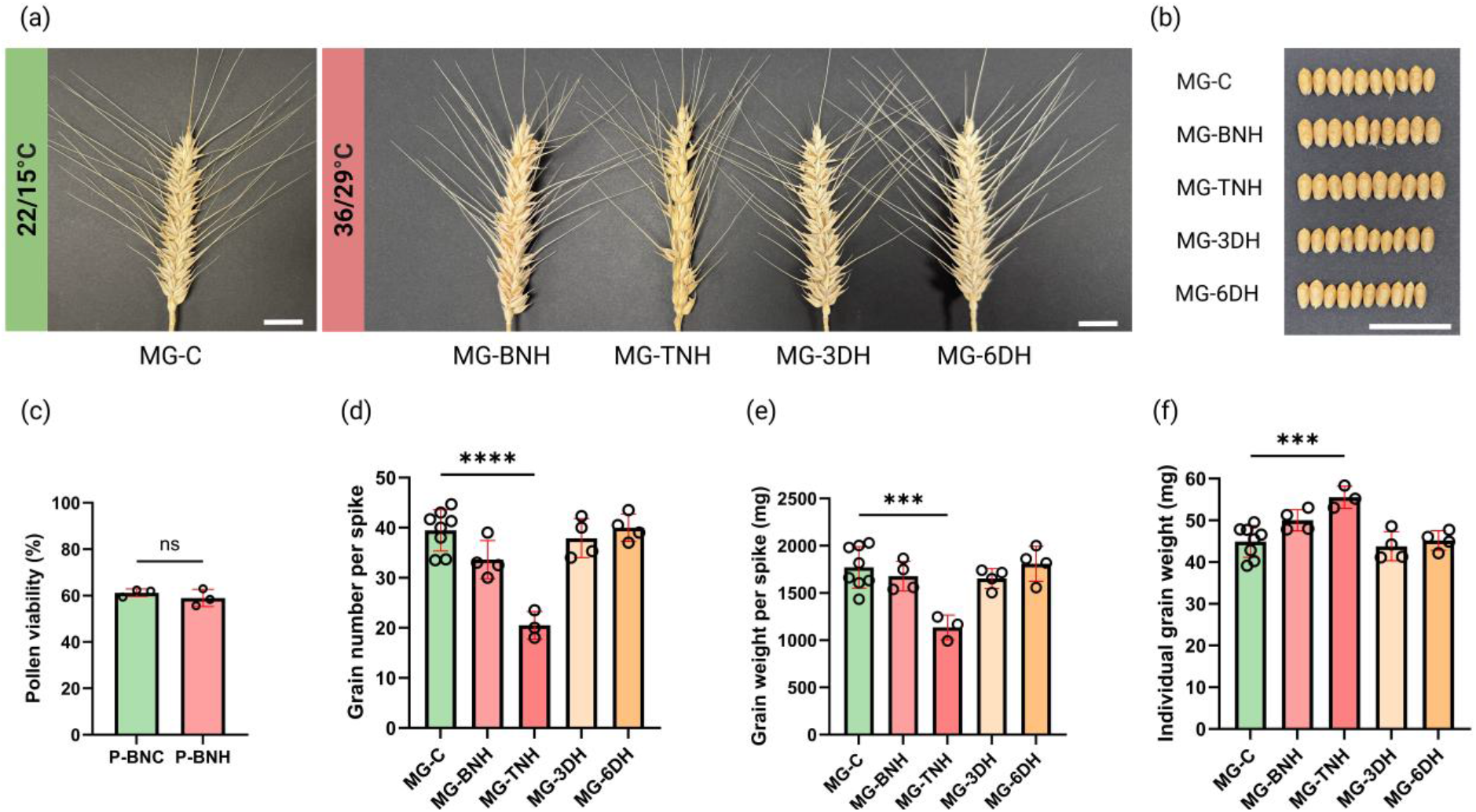
Effects of short-term heat stress (36/29 ℃ for 48 h) at defined developmental stages on plant reproductive and yield-related traits. (a) Representative spikes from control plants (MG-C) and plants heated at the binucleate (MG-BNH), trinucleate (MG-TNH), 3 days after anthesis (MG-3DH), or 6 days after anthesis (MG-6DH) stages. Scale bar: 2 cm. (b) Representative grains from each treatment. Scale bar: 2 cm. (c) Pollen viability (%) of mature pollen from control (P-BNC) and heat-treated plants at the binucleate stage (P-BNH). (d–f) Yield components measured in mature grains: grain number per spike (d), grain weight per spike (e), and individual grain weight (f). Data represent mean ± SE (n ≥ 3 biological replicates), with individual values shown. Statistical significance was assessed using unpaired t-tests or one-way ANOVA (\**p* < 0.05; \*\**p* < 0.01; \*\*\**p* < 0.001; \*\*\*\**p* < 0.0001; ns, not significant).

### Heat does *not* impair pollen viability but increases floret abortion

Exposure to 36/29°C for 48 h during BN did not significantly reduce pollen viability at anthesis (Fig. 2c), indicating that pollen heated during the BN stage remains relatively tolerant to elevated temperatures. Despite the maintenance of pollen viability, heat at BN or TN substantially increased floret abortion on primary tillers. When primary or secondary florets aborted, the third (and occasionally fourth) floret within the spikelet continued to develop, although it was developmentally less advanced. Under control conditions, spikelets typically supported the development of up to three florets. Together, these findings suggest potential sensitivity of female reproductive tissues (e.g. ovule or ovary function, double fertilisation or early grain initiation) which may be disrupted by short-term heat at BN and TN, leading to increased reproductive failure even when the pollen remains viable.

### Heat during anthesis drives the largest reduction in grain set

Heat exposure had stage-specific impacts on yield components. Grain number per spike decreased by ∼15% at BN (*ns*) and ∼48% at TN (*p* < 0.0001), whereas 3 DAA and 6 DAA were comparable to the control (Fig. 2d). Total grain weight per spike broadly mirrored this pattern (Fig. 2e): the strongest reduction at TN (∼36%, *p* < 0.001) and a modest decrease at BN, while 3 DAA and 6 DAA were largely similar to the control. In contrast, individual grain weight tended to increase at BN (∼11%, *ns*) and TN (∼24%, *p* < 0.001) and was similar to controls after heat at 3 DAA and 6 DAA (Fig. 2f). Collectively, these results indicate that anthesis represents a critical developmental window linking transient stress exposure to irreversible yield loss, whereas pre-(BN) and post-anthesis (3 DAA and 6 DAA) heat has minimal effects on yield.

### Heat at anthesis strongly alters the amino acid composition in mature grain

Amino acid composition of milled grain showed stage-dependent changes under heat (Fig. 3). Most amino acids (n = 11) displayed consistent decreases in relative amino acid composition when heat was applied at the trinucleate stage (MG-TNH), with more moderate or smaller changes at MG-BNH, MG-3DH, and MG-6DH. Essential amino acids, including histidine, leucine, lysine, methionine, threonine and valine, were most strongly reduced at MG-TNH, followed by MG-3DH, MG-BNH and MG-6DH, with only some showing intermediate reductions. In contrast, phenylalanine increased at MG-TNH, with no significant changes at the other stages, whereas isoleucine remained unchanged across all treatments. Proline (most prominently at MG-TNH) and glutamic acid (across all stages) increased under heat, with tyrosine also showing a non-significant upward trend. Other amino acids, including alanine, arginine, aspartic acid, glycine and serine, exhibited their largest reductions relative to the control at MG-TNH, followed by MG-3DH, MG-BNH, and MG-6DH, with only moderate shifts at some stages. Overall, post-meiotic heat stress altered amino acid metabolism during grain development, with anthesis (TN) being the most sensitive stage and 6 DAA the least affected.

**Figure 3.**
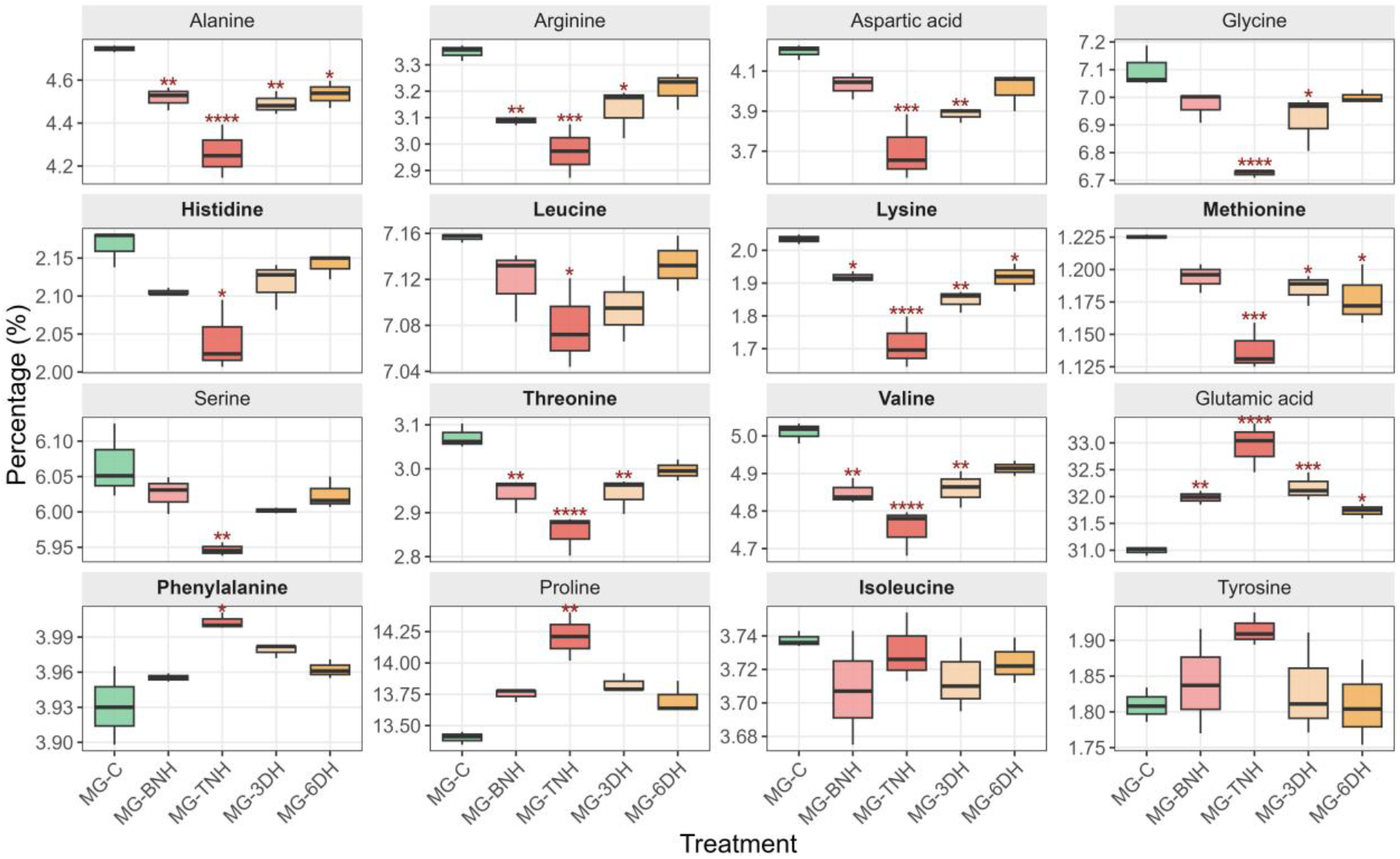
Relative amino acid composition of whole-grain flour (70 µm) under control and heat treatments (36/29°C for 48 h). Boxplots show the proportion of each amino acid, expressed as a percentage of the total measured amino acid pool, across MG-C, MG-BNH, MG-TNH, MG-3DH, and MG-6DH (n = 3 biological replicates per treatment). Asterisks indicate significant differences between each heat treatment and the control (MG-C). Early heat (M-BNH and M-TNH) produced the strongest compositional shifts, including reductions in most amino acids and increases in glutamic acid, phenylalanine, and proline, while later heat treatments during the early grain-filling stages (MG-3DH and MG-6DH) showed smaller or nonsignificant changes. Statistical comparisons versus MG-C were performed using one-way ANOVA with Dunnett’s test or Kruskal–Wallis with Dunn’s test when assumptions were not met. Essential amino acids are shown in bold. Significance levels: \**p* < 0.05; \*\**p* < 0.01; \*\*\**p* < 0.001; \*\*\*\**p* < 0.0001.

### NIR spectroscopy reveals moderate treatment-dependent variation in mature grain

PCA of the NIR spectra showed partial separation among treatments, with PC1 (85.1% of variance) capturing the dominant variation and PC2 (7.7%) contributing additional structure (Fig. S2). Treatments exhibited overlapping distributions; however, distinct clustering was observed for MG-TNH and MG-6DH samples along PC1 and PC2, indicating consistent spectral differences under these conditions. In contrast, MG-C, MG-BNH and MG-3DH samples displayed greater dispersion, reflecting higher within-group variability. Overall, the NIR spectra captured moderate treatment effects, although separation among all developmental stages remained incomplete.

### Micro-CT analysis reveals heat-induced alteration of crease architecture

Based on morphological assessment, the crease region of the mature grain was selected for quantitative micro-CT analysis because it exhibited the most consistent and reproducible structural differences between control and heat-treated samples (Fig. 4a). Two complementary geometric parameters were quantified from central transverse micro-CT sections (Fig. 4b). MG-TNH exhibited a significantly increased CD (∼67%) compared with MG-C (*p* < 0.05; Fig. 4c). This indicates deeper penetration of the crease cavity into the grain interior following heat stress at the trinucleate stage. When measured at 50% of CD, CW was reduced by ∼86% in MG-TNH relative to MG-C; however, this difference was not statistically significant (*p* < 0.06; Fig. 4d), consistent with increased variability associated with cavity geometry. In contrast, MG-BNH did not differ significantly from the control for either CD or CW. Overall, these data demonstrate that heat stress at TN specifically alters crease architecture, whereas exposure at BN does not produce equivalent structural changes. This stage specificity suggests that heat stress at the trinucleate stage is the primary driver of the observed structural reorganisation.

**Figure 4.**
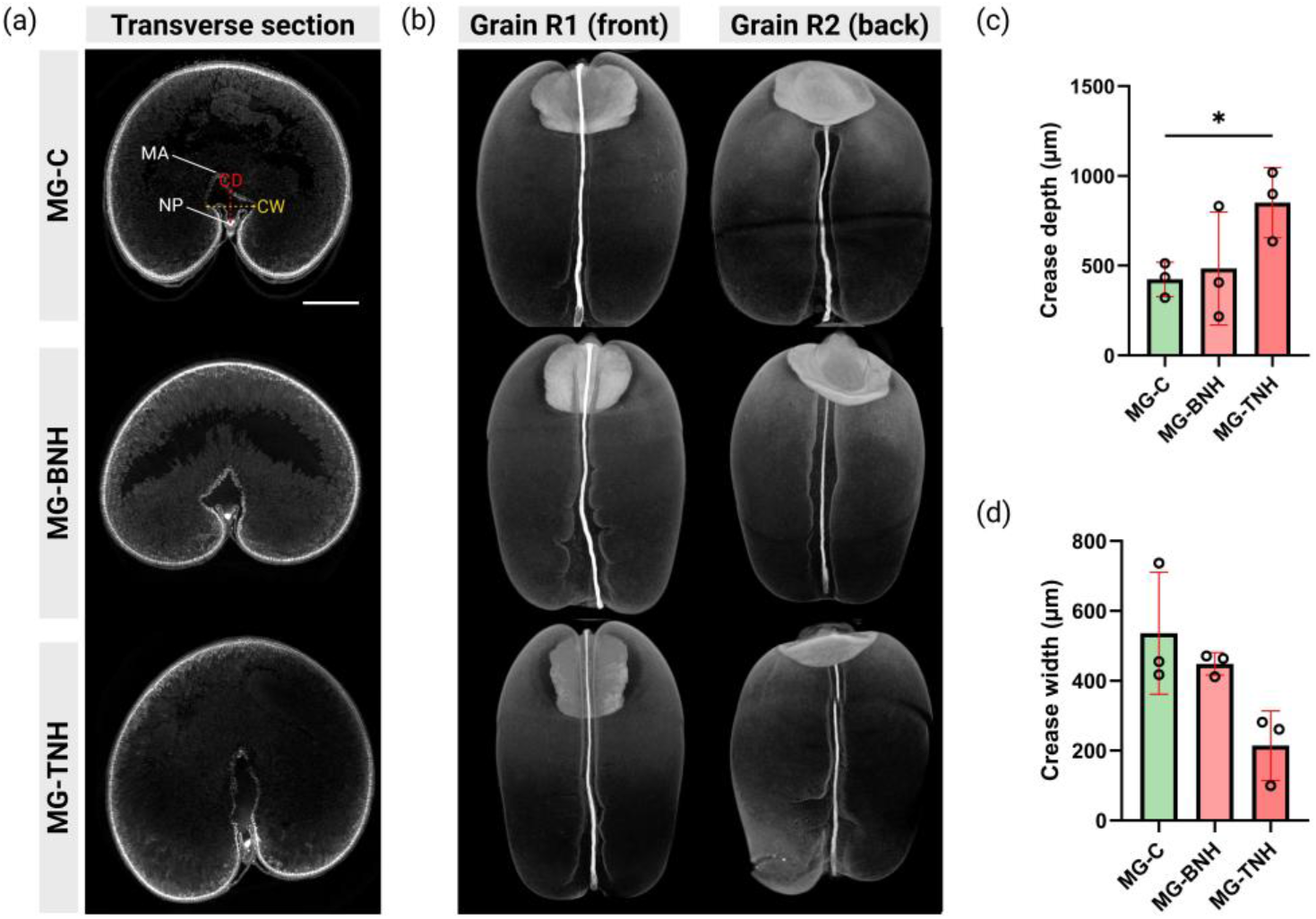
Heat stress (36/29°C for 48 h) at the trinucleate stage alters crease architecture. (a) Transverse micro-CT sections of MG-C, MG-BNH and MG-TNH grains. MA, modified aleurone; NP, nucellar projection; CD, crease depth; CW, crease width measured at 50% of CD along the crease axis. Scale bar = 500 μm. (b) Front and back 3D micro-CT reconstruction of representative MG-C, MG-BNH, and MG-TNH grains. MG-C grains show smooth crease margins, whereas MG-BNH and MG-TNH grains exhibit localised groove-like surface deformations adjacent to the crease. (c) Quantification of crease depth (CD), showing a significant increase in MG-TNH compared with MG-C. (d) Quantification of crease width (CW) measured at 50% of CD along the crease axis. CW shows an apparent reduction in MG-TNH relative to MG-C, although this difference was not statistically significant. Statistical comparisons versus MG-C were performed using unpaired two-tailed *t*-tests with Welch’s correction. Significance level: \**p* < 0.05.

To provide spatial context for the quantitative differences observed in transverse sections, three-dimensional (3D) micro-CT reconstructions were generated for representative grains from MG-C, MG-BNH, and MG-TNH. Front and back views of two independent grains per treatment illustrate that the alterations detected in transverse sections extend along the length of the crease. Compared with the smooth crease margins observed in MG-C grains, both MG-BNH and MG-TNH grains exhibit localised groove-like surface deformations in the vicinity of the crease. These features are evident on both faces of the grains and are more pronounced in MG-TNH, consistent with the increased crease penetration quantified in trinucleate heat-stressed samples. Together, the 3D reconstructions qualitatively reinforce the stage-specific structural reorganisation of the crease identified by cross-sectional analysis.

### Heat-induced epigenetic reprogramming is stage-dependent

Global methylation levels were highly similar among IMG-C, IMG-BNH, and IMG-TNH, indicating that heat stress did *not* induce widespread epigenomic reprogramming (Fig. S3a). Instead, heat responses were manifested as differentially methylated regions (DMRs), predominantly in the CG context captured by RRBS, distributed unevenly across chromosomes (Fig. S3b). In spite of producing less pronounced phenotypic effects than IMG-TNH, IMG-BNH exhibited a more extensive methylation response (Fig. 5a; Supplementary Data S1). This difference was particularly evident in promoter-proximal regions, where IMG-BNH contained more than double the number of DMRs observed in IMG-TNH (Fig. 5b), highlighting enhanced targeting of *cis*-regulatory regions during the binucleate stage. Interestingly, only a small proportion of differentially methylated genes (DMGs) was shared between the two stages, indicating that heat induces strongly stage-specific epigenetic responses (Fig. 5d–e).

**Figure 5.**
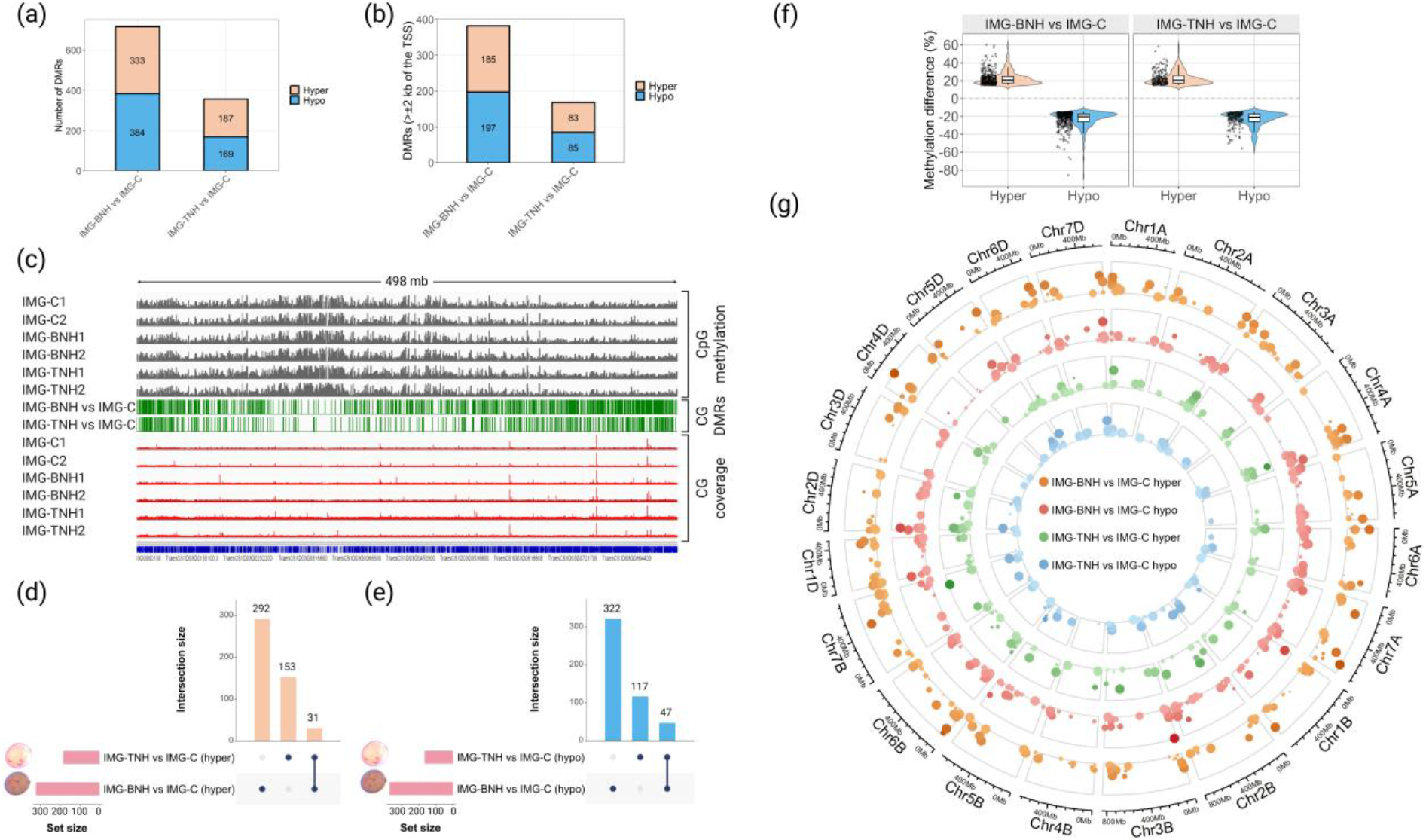
Genome-wide and locus-specific DNA methylation responses of developing grains exposed to heat (36/29°C for 48 h) during the binucleate and trinucleate stages. (a) Number of DMRs for each comparison, with IMG-BNH vs IMG-C containing a greater number of hyper- and hypo-DMRs compared with IMG-TNH vs IMG-C. (b) Number of promoter-proximal (> ±2 kb of the TSS) hyper- and hypo-DMRs for each comparison. (c) Whole-chromosome CG methylation visualisation for Chromosome 1D, selected as the representative example. Tracks show CG methylation (grey) for all samples, DMRs (green) for the two comparisons, and CG coverage (red) for all samples. (d–e) UpSet plots showing intersections of hyper-DMGs (d) and hypo-DMGs (Conway et al. 2017) (e) among IMG-BNH vs IMG-C and IMG-TNH vs IMG-C, highlighting strong stage-specific epigenetic responses. (f) Effect-size distributions of hyper- and hypo-DMRs. Violin and boxplots show methylation differences (%) for significant DMRs, illustrating more extreme hypo-methylation relative to hyper-methylation. (g) Circos map of hyper- and hypo-DMRs (≥15% methylation difference, *q*≤0.01). Rings (outer→inner) represent IMG-BNH hyper, IMG-BNH hypo, IMG-TNH hyper, and IMG-TNH hypo DMRs. Darker points indicate greater methylation difference, while larger points indicate stronger statistical support.

DMRs were unevenly distributed across A, B, and D subgenomes, with each developmental stage showing distinct chromosomal hotspots (Fig. 5c; Fig. S3b). Across both stages, demethylation events (down to –85.5% in IMG-BNH and –55.7% in IMG-TNH) were generally larger in magnitude than methylation gains (up to +60.1% in IMG-BNH and +58.1% in IMG-TNH) (Fig. 5f), consistent with stronger demethylation caused by heat, particularly at the binucleate stage. Highly hypo-methylated loci included genes encoding an AB hydrolase-1 domain-containing protein (*TraesCS1D03G0003300*), 3-ketoacyl-CoA synthase (*TraesCS7D03G1226000*), and a hyperosmolality-gated Ca^2+^ permeable channel (*TraesCS1B03G0073000*)—genes collectively involved in lipid remodelling, very-long-chain fatty acid biosynthesis, and Ca²⁺-mediated stress signalling—as well as several uncharacterised genes (*TraesCS5B03G1252800LC*, *TraesCS2B03G1219100LC*, *TraesCS2D03G0135000LC* and *TraesCS3B03G0112100LC*). IMG-BNH showed a higher density of DMRs, larger effect sizes and stronger statistical support than IMG-TNH. Although a substantial proportion of DMRs overlapped between stages, IMG-BNH consistently exhibited more pronounced methylation changes and occasional opposing methylation responses between stages, indicating developmentally distinct regulatory responses (Fig. 5g).

Collectively, IMG-BNH exhibited broader and stronger methylation responses, particularly at promoter-proximal regions, whereas TNH displayed a more targeted and attenuated pattern. These findings indicate that BN exhibits greater epigenetic responsiveness, potentially enabling transcriptional reprogramming prior to fertilisation. However, this increased molecular responsiveness does not cause stronger phenotypic effects.

### Spatial RNA-seq resolves tissue-specific transcriptional domains in developing grains

Spatial transcriptomic analysis resolved coherent transcriptional domains that corresponded closely to major anatomical regions of the developing grain, including the scutellum, plumule, nucellar projection (NP), vascular tissue, modified aleurone and starchy endosperm-aleurone interface (Fig. 6a). Marker gene analysis identified distinct transcriptional signatures associated with individual clusters (Fig. 6b), providing support for cluster annotation and facilitating assignment of biologically meaningful tissue identities. Heat stress primarily altered transcriptional activity *within* existing spatial domains rather than reorganising gene expression patterns across tissues.

**Figure 6.**
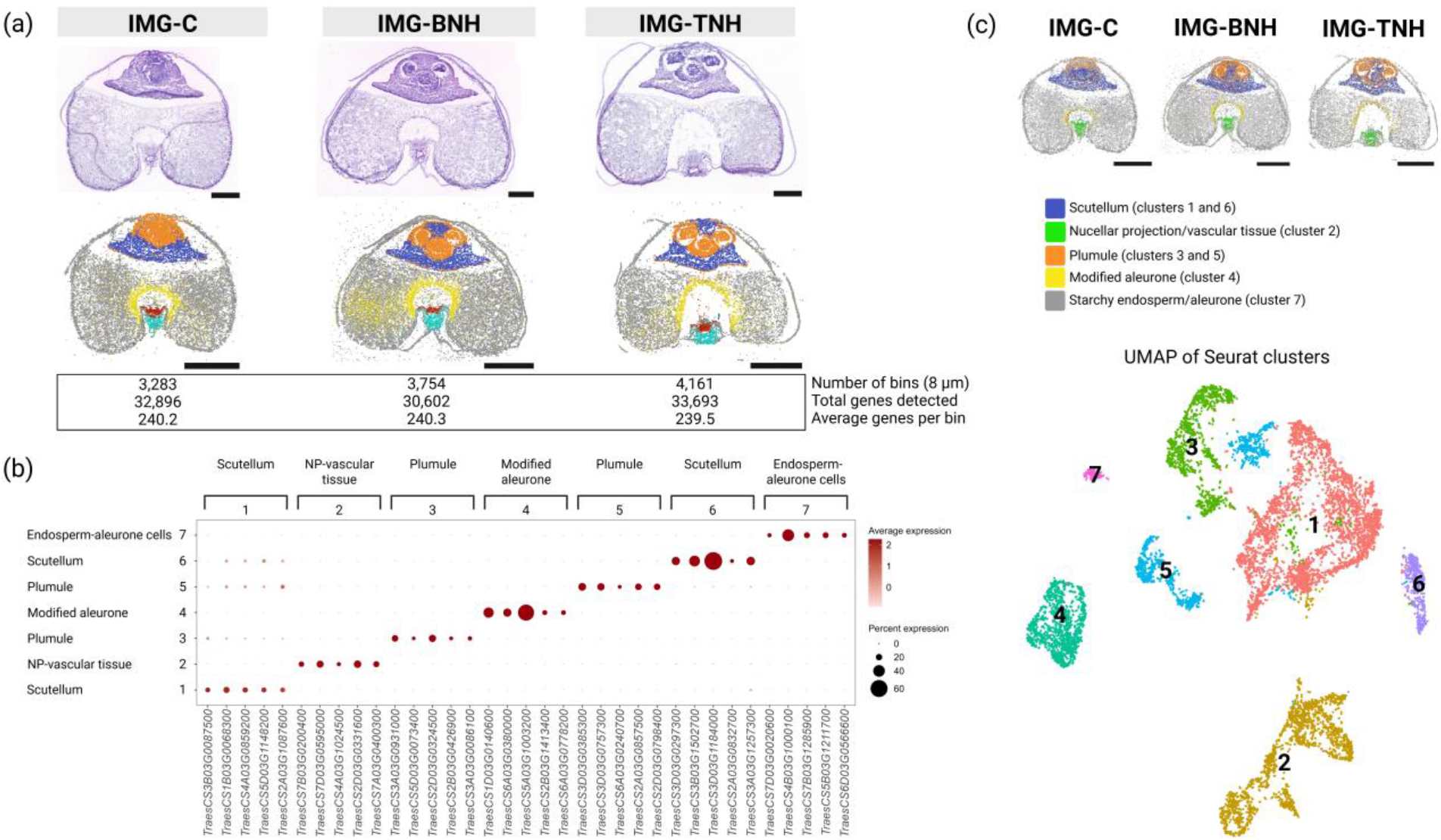
Spatial organisation and transcriptional clustering of developing wheat grains under control and heat stress (36/29°C for 48 h). (a) H&E-stained transverse sections of developing wheat grains from control (IMG-C), binucleate heat-treated (IMG-BNH), and trinucleate heat-treated (IMG-TNH) conditions, with corresponding spatial transcriptomic visualisation showing global gene expression patterns across tissue sections. Spatial patterns reveal coherent anatomical regions consistent with underlying histology. (b) Dot plot showing the five highest-ranking marker genes identified for each Seurat cluster. Dot size represents the percentage of spatial bins within a cluster in which a gene was detected, while dot colour represents the average scaled expression level. Marker genes were selected based on average log₂ fold change and statistical significance. (c) Unsupervised clustering of spatial transcriptomic profiles using Seurat resolves seven transcriptional clusters, shown in UMAP space (bottom). Cluster identities were assigned based on marker genes identified using *FindMarkers* and correspond to biologically meaningful tissue regions. Spatial projection of annotated clusters (top) reveals correspondence with anatomical structures, including the scutellum (clusters 1 and 6), nucellar projection/vascular-associated region (cluster 2), plumule (clusters 3 and 5), modified aleurone (cluster 4), and starchy endosperm–aleurone interface (cluster 7). Scale bars: 0.5 mm (H&E), 1 mm (spatial plots).

Unsupervised clustering analysis identified seven transcriptionally distinct clusters, corresponding to biologically meaningful tissue compartments, confirming that spatial transcriptomics robustly captures tissue-specific gene expression programs in developing wheat grain (Fig. 6c; Fig. S4). These clusters exhibited partial spatial overlap in embryo-associated regions (scutellum and plumule), indicating shared transcriptional responses within common anatomical compartments. Together, these data demonstrate that heat responses are spatially organised and constrained by tissue identity, generating stage-dependent molecular signatures within discrete grain tissues rather than uniform transcriptional changes across the grain.

### Spatial RNA-seq reveals stage-dependent molecular imprints of pre-anthesis and anthesis heat stress on grain development

Pseudobulk differential expression analysis revealed markedly stronger transcriptional responses following heat exposure at the binucleate stage than at the trinucleate stage (Fig. 7a; Supplementary Data S2). Most differentially expressed genes (DEGs) were stage-specific, with only limited overlap between IMG-BNH and IMG-TNH, demonstrating that closely spaced developmental stages generate distinct transcriptional signatures (Fig. 7b–c).

**Figure 7.**
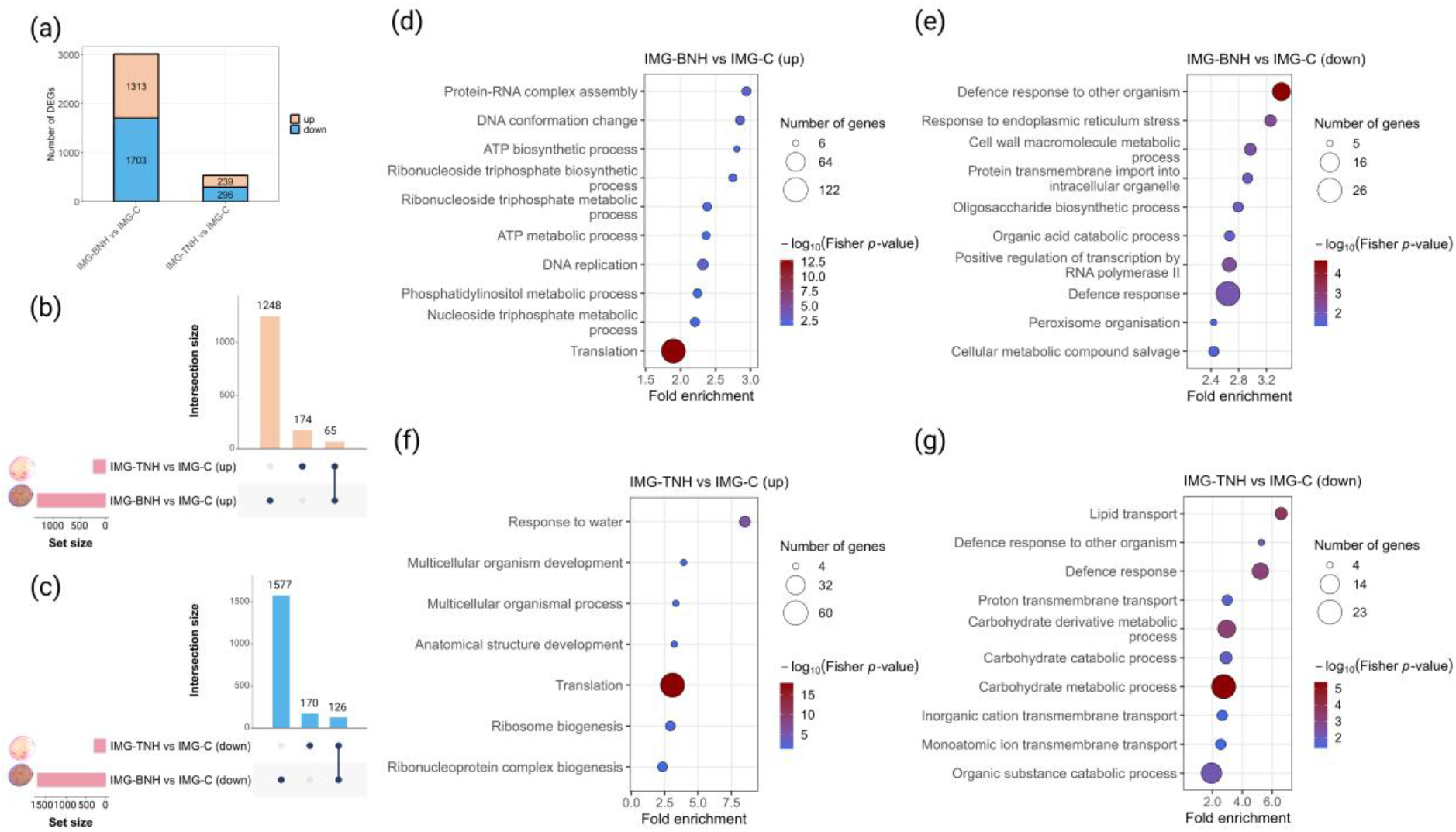
Pseudobulk differential expression and functional enrichment of spatial transcriptomes under control and heat-stress (36/29°C for 48 h) conditions. (a) Number of DEGs identified by pseudobulk differential expression analysis in IMG-BNH vs IMG-C and IMG-TNH vs IMG-C, showing a substantially stronger response in IMG-BNH. (b–c) UpSet plots showing intersections of up-regulated (b) and down-regulated (c) genes, highlighting predominantly stage-specific transcriptional responses to heat. (d–g) Bubble plots showing the top enriched GO terms within the Biological Process (BP) category among up- and down-regulated genes in IMG-BNH vs IMG-C (d–e) and IMG-TNH vs IMG-C (f–g). GO enrichment was performed with topGO using the *elim* algorithm, with all expressed genes in the corresponding treatment as the background. Bubble size represents the number of contributing genes, and colour indicates −log_10_ (Fisher *p*-value).

Gene ontology (GO) analysis revealed functionally distinct transcriptional responses between stages. Heat exposure at BN promoted pathways associated with translation, ribosome function, DNA maintenance and replication, and energy metabolism, while suppressing defence responses, endoplasmic reticulum (ER)-associated proteostasis and transcriptional regulation (Fig. 7d–e). In contrast, heat stress at TN elicited a more restricted transcriptional reprogramming characterised by activation of developmental processes and suppression of carbohydrate metabolism, transport and sink-related functions (Fig. 7f–g).

Comparison of differential methylation and gene expression revealed no significant genome-wide correlation between methylation and transcriptional responses (Spearman *R* = –0.081 to 0.043, *p* > 0.05) (Fig. S5), indicating that transcriptional reprogramming is determined primarily by gene-specific and stage-dependent regulatory mechanisms rather than global methylation patterns.

### Spatial RNA-seq reveals tissue-specific transcriptional reprogramming under heat stress

Spatial expression of representative marker genes corresponded closely to known seed anatomy, confirming accurate spatial resolution of major tissue domains (Fig. 8a). At the whole-grain level, earlier heat events activated transcriptional programs associated with translation, DNA maintenance and replication, and energy metabolism, while suppressing defence responses, ER-associated proteostasis, transcriptional regulation, carbohydrate metabolism and transport (Fig. 8b). These responses were more pronounced in IMG-BNH than IMG-TNH, indicating stronger transcriptional reprogramming following binucleate-stage heat stress.

**Figure 8.**
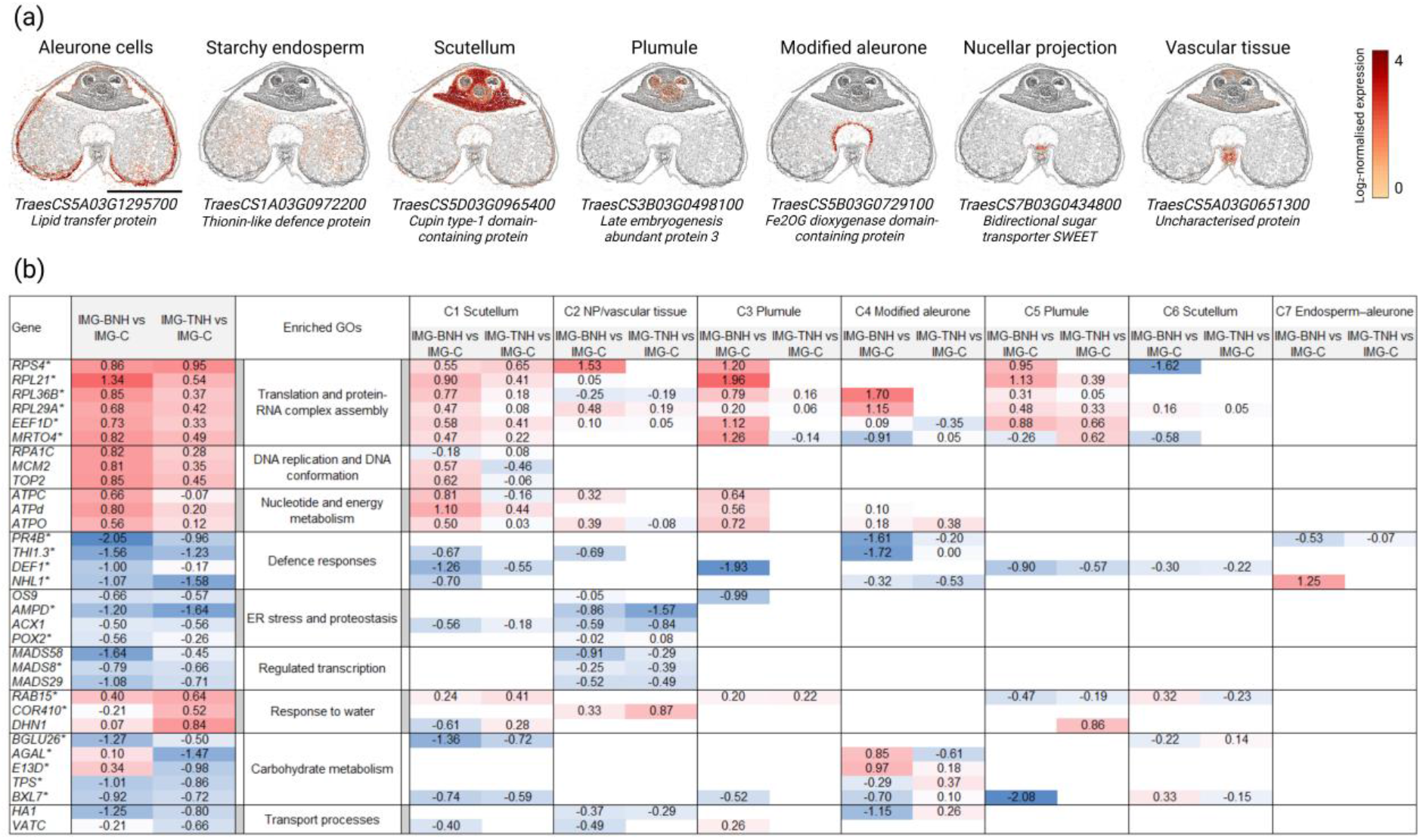
Representative gene-level heatmap of enriched biological processes under heat stress (36/29°C for 48 h). (a) Representative spatial anchor genes showing distinct localisation patterns corresponding to major tissue and interface domains. (b) Heatmap of representative genes selected from significantly enriched GO categories, based on functional annotation and consistent regulation across conditions, identified in IMG-BNH and IMG-TNH relative to the control. Genes were curated based on functional relevance and transcriptional behaviour to highlight dominant biological processes rather than exhaustive GO membership. Log₂ fold changes are shown for IMG-BNH and IMG-TNH, with red indicating up-regulation and blue indicating down-regulation. A fixed, symmetric colour scale centred at zero was applied across conditions to enable direct comparison. To assess spatial modulation, the same gene set was evaluated across all clusters. While the direction of regulation for major biological processes was broadly conserved, differences in magnitude revealed cluster-specific modulation of shared transcriptional programs. Some representative genes showed limited detection in specific clusters, reflecting restricted or heterogeneous expression at the tissue-domain level rather than absence of regulation in whole-sample analyses. Cluster-marker genes are indicated by an asterisk.

Spatially, heat-responsive programs were conserved across tissue domains but varied markedly in magnitude. Embryo-associated tissues, including the scutellum (C1) and plumule (C3, C5), showed the strongest activation of biosynthetic processes. Translation-associated genes such as *RPS4* (40S ribosomal protein S4), *RPL21* (60S ribosomal protein L21-1-like) and *EEF1D* (Elongation factor 1-delta 1) were strongly induced in these regions, whereas modified aleurone (C4) and NP/vascular tissues (C2) exhibited weaker responses and localised suppression of biosynthetic activity (Fig. 8b).

Defence responses were spatially heterogeneous, with defence markers *PR4B* (wheatwin-2), *THI1.3* (purothionin A-1) and *DEF1* (defensin-like protein 1) showing the strongest suppression in modified aleurone (C4) and plumule tissues (C3). Similarly, ER-associated proteostasis and regulated transcription were predominantly repressed in NP/vascular tissues (C2), exemplified by the down-regulation of *ACX1* (peroxisomal acyl-coenzyme A oxidase 1-like) and *OS9* (protein OS-9 homolog), and the developmental regulators *MADS58* and *MADS29* (Fig. 8b).

‘Carbohydrate metabolism’ was preferentially suppressed in embryo-associated tissues, exemplified by down-regulation of the carbohydrate-remodelling genes *BGLU26* (beta-glucosidase 26-like) and *BXL7* (probable beta-D-xylosidase 7 isoform X2), indicating reduced carbon utilisation within the developing embryo. Additional carbohydrate-related genes, including *AGAL* (alpha-galactosidase) and *TPS* (trehalose-6-phosphate synthase), exhibited highly localised responses within modified aleurone tissues, indicating spatial partitioning of carbohydrate metabolism across distinct grain domains. Transport-associated pathways, exemplified by *HA1* (plasma membrane ATPase isoform X2) and *VATC* (V-type proton ATPase subunit C-like isoform X1), were predominantly repressed in modified aleurone and NP/vascular tissues, indicating localised disruption of assimilate transport and sink activity within the crease region. In contrast, osmotic stress responses were activated in a tissue-specific manner, with the dehydrin genes *COR410*, *DHN1* and *RAB15* showing strong induction in embryo-associated and NP/vascular tissues (Fig. 8b).

Overall, these results demonstrate that heat stress induces a broadly conserved transcriptional response across developing wheat grains, but that the magnitude and spatial distribution of these responses are strongly constrained by tissue identity. Modified aleurone, nucellar projection and vascular tissues represent major sites of defence, proteostasis and transcriptional suppression, whereas genes associated with biosynthetic pathways were linked with embryo-associated tissues. These effects were most pronounced following binucleate-stage heat exposure, whereas responses at the trinucleate stage were more moderate but retained a similar spatial organisation.

### Spatial RNA-seq reveals expansion and amplification of heat-responsive domains

Spatial stress scoring revealed a progressive increase in heat-responsive transcriptional activity from control to IMG-BNH and IMG-TNH (Fig. 9a–b). While global stress responses were broadly distributed across developing grains, marker-derived stress responses were concentrated within their defined tissue regions and were highly reproducible across biological replicates (Fig. S6). Both stress metrics differed significantly across treatments (Kruskal–Wallis test, *p* < 2.2 × 10⁻¹⁶).

**Figure 9.**
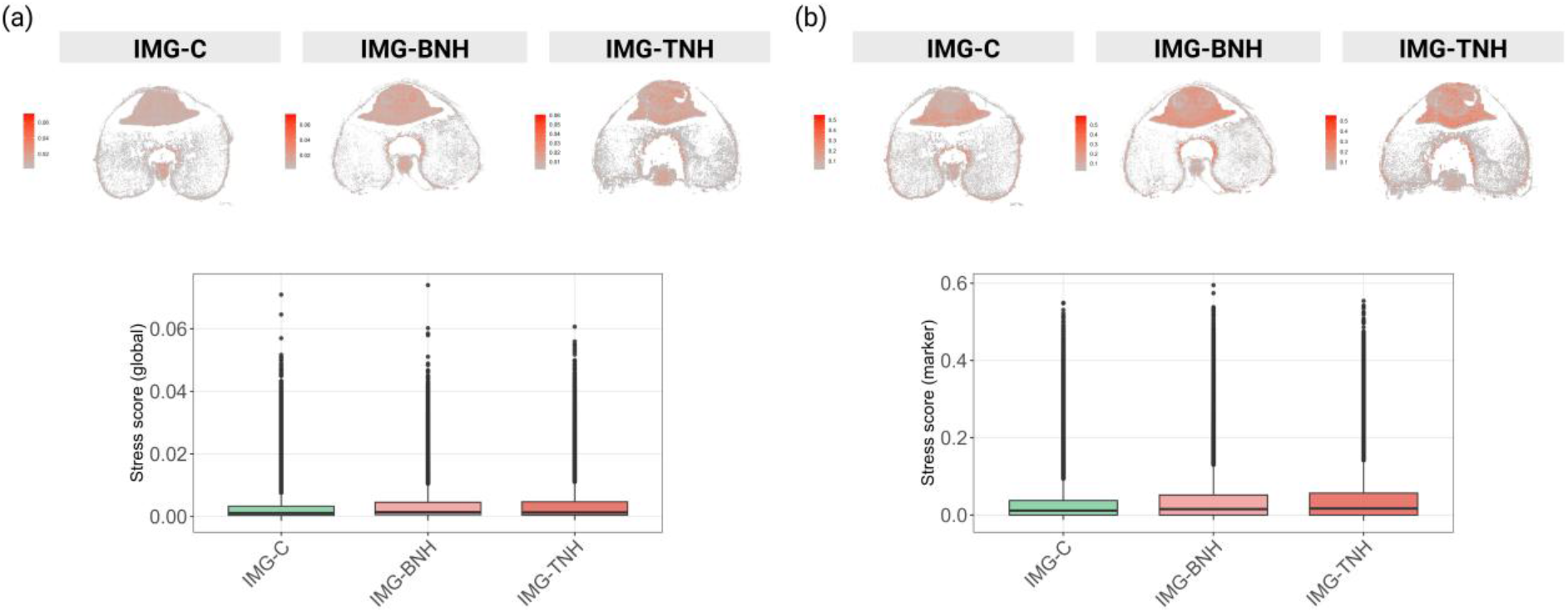
Global and marker-based heat-response gene programs. To assess spatial transcriptional responses to heat stress (36/29°C for 48 h), we generated two stress scores (calculated for each spatial bin using Seurat’s *AddModuleScore*): a global stress score based on 14,585 DEGs and a marker-derived stress score based on 697 treatment- and cluster-associated marker genes. (a) Spatial distribution of the global-derived stress-response program across representative sections from control (IMG-C), IMG-BNH and IMG-TNH samples. The global stress score shows a broadly distributed transcriptional response across the tissue, with magnitude increasing progressively from control to IMG-TNH. (b) Spatial distribution of the marker-derived stress-response program. Elevated values are more strongly concentrated within specific tissue domains. Boxplots below each panel show the distribution of program scores across all spatial bins and biological replicates. Each point represents an individual spatial bin.

The proportion of high-activity spatial bins increased from 7.2% in control samples to 11.4% and 11.9% in IMG-BNH and IMG-TNH, respectively (χ² = 4118.9, *p* < 2.2 × 10⁻¹⁶), illustrating expansion of stress-responsive domains following heat exposure. While both IMG-BNH and IMG-TNH recruited a larger proportion of tissue into the transcriptional response than the control, domain extent differed only marginally between IMG-BNH and IMG-TNH. In contrast, median, upper-quartile, and 90^th^ percentile marker-derived stress scores increased progressively from control to IMG-BNH and IMG-TNH, indicating stronger transcriptional activation within stress-responsive domains.

Together, these results demonstrate that heat stress both expands the spatial footprint of responsive tissues and amplifies transcriptional activity within those domains. Anthesis heat stress was characterised by elevated stress-score distributions despite only a modest increase in domain extent relative to IMG-BNH, suggesting that stronger heat responses are driven primarily by increased transcriptional activity within stress-responsive regions rather than substantial additional domain expansion.

### Heat stress induces coordinated metabolic reprogramming and network-level reorganisation

To investigate metabolic responses to heat stress during grain development, a curated set of metabolites was grouped into functional categories, including RNA metabolism, nucleotide metabolism, central carbon metabolism, secondary metabolism, amino acid metabolism, methylation-related metabolites, redox metabolism, and stress-associated metabolites. Heat stress increased metabolites associated with nucleotide signalling and flavonoid-rich secondary metabolism, while reducing the abundance of amino acid-related metabolites (Fig. 10a).

**Figure 10.**
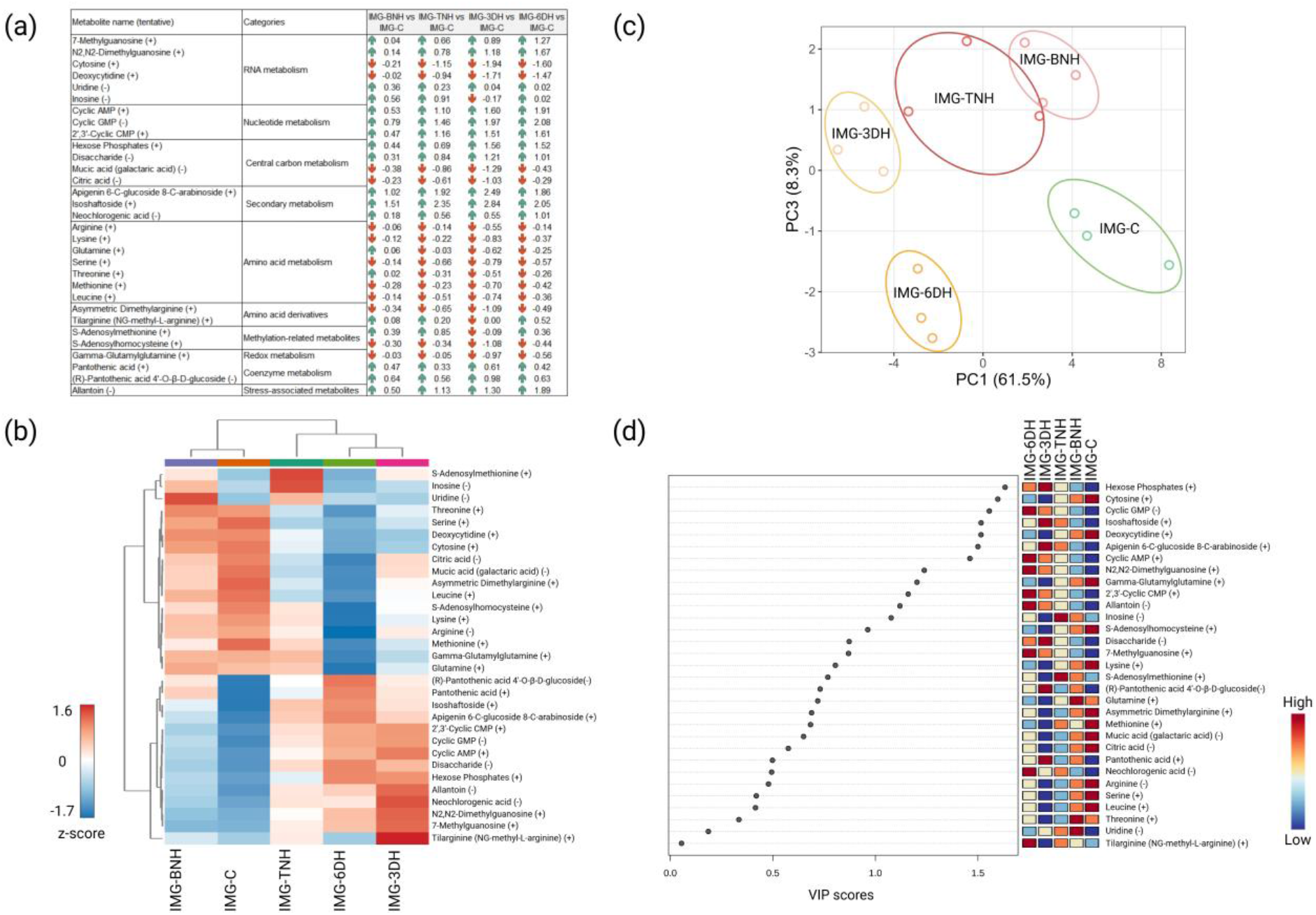
Integrated metabolomic analysis under heat stress. (a) Selected metabolites showing differential abundance under heat stress (36/29°C for 48 h) (IMG-BNH, IMG-TNH, IMG-3DH and IMG-6DH) relative to control (IMG-C). Values represent fold changes, with arrows indicating direction of change (↑ increase, ↓ decrease). Metabolites are grouped by functional categories. The symbols (+) and (−) indicate detection in positive and negative ion modes, respectively. (b) Heatmap of log2-transformed metabolite abundances after z-score scaling, illustrating relative changes across conditions. (c) Principal component analysis (PCA) score plot based on log2-transformed and scaled metabolite data (z-score scaling). (d) Variable importance in projection (VIP) scores derived from sparse PLS-DA identifying metabolites contributing to multivariate separation.

Hierarchical clustering revealed clear separation between pre-anthesis (IMG-BNH) and post-anthesis (IMG-3DH and IMG-6DH) heat treatments, with IMG-TNH exhibiting a distinct metabolic profile (Fig. 10b). Metabolites associated with nucleotide signalling (e.g. cAMP and cGMP), modified nucleosides (e.g. 7-methylguanosine and N2,N2-dimethylguanosine), and flavonoid-rich secondary metabolism (e.g. isoschaftoside and apigenin derivatives) were consistently elevated under heat stress. In contrast, amino acid metabolism and several central carbon intermediates, including citric acid and mucic acid, were reduced or remained comparatively stable, indicating a shift away from primary metabolism. The stress-associated metabolite allantoin was strongly enriched under heat stress, whereas nucleotide metabolites (e.g. inosine, uridine), methylation-related metabolites (e.g. S-adenosylmethionine and S-adenosylhomocysteine) and the redox-related metabolite gamma-glutamylglutamine generally declined under heat stress.

PCA and sparse partial least squares discriminant analysis (sPLS-DA) confirmed clear separation between heat-stressed (IMG-BNH, IMG-TNH, IMG-3DH and IMG-6DH) and control (IMG-C) samples and identified carbohydrate-related intermediates, cyclic nucleotides, flavonoid derivatives and modified nucleosides as major contributors to treatment differentiation (Fig. 10c–d). The distinct positioning of IMG-BNH, IMG-TNH, IMG-3DH and IMG-6DH indicates that heat stress generates stage-specific metabolic states that persist into grain development.

Together, these analyses demonstrate that heat stress induces coordinated metabolome-wide reorganisation, characterised by a shift from primary metabolic processes toward regulatory and stress-associated pathways, including nucleotide signalling, modified nucleosides, and secondary metabolite biosynthesis.

### DESI-MSI reveals tissue-specific localisation of lipid and flavonoid ions across treatments

Target metabolites were detected at relatively low signal intensity, consistent with the reduced ionisation efficiency of polar compounds in DESI-MSI. However, higher-abundance ions, including lipid- and flavonoid-related species, exhibited distinct and reproducible spatial distributions across grain tissues (Fig. 11). Selected ions in the *m/z* 575–617 range, including m/z ∼617.49, were consistently enriched in the embryo and aleurone regions, whereas ions such as *m/z* ∼543.13 were predominantly localised to the endosperm. In contrast, ions at *m/z* ∼365.11 and ∼381.09 were associated with the aleurone layer, modified aleurone and crease regions. These spatial patterns were highly consistent across independent replicate sections for all treatments.

**Figure 11.**
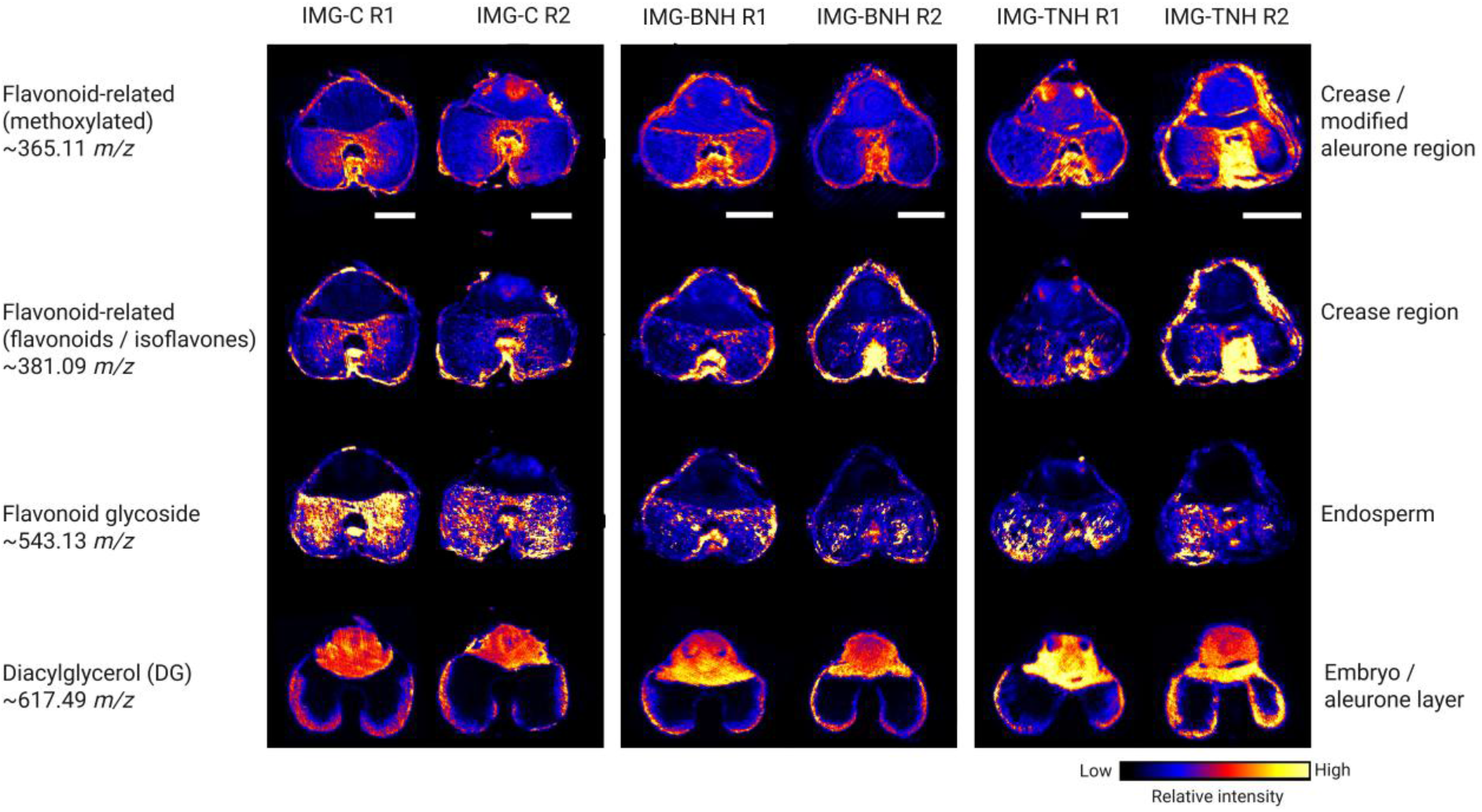
DESI-MSI of wheat grain transverse sections showing spatial distribution of selected ions (*m/z* ∼365.11, ∼381.09, ∼543.13 and ∼617.5) across IMG-C, IMG-BNH and IMG-TNH treatments. Ion images were visualised using a shared intensity scale to enable qualitative comparison across sections. Major anatomical regions including embryo, aleurone layer, endosperm, crease and modified aleurone are indicated. Putative ion annotation was performed using the LIPID MAPS database with a strict mass tolerance (±0.01 Da), and all selected ions showed the most consistent matches under sodium-adduct conditions ([M+Na]+). Due to the presence of multiple structural isomers and the absence of MS/MS confirmation, annotations are reported at the compound-class level. Scale bar = 1 mm.

Putative annotation of spatially resolved ions indicated flavonoid-related metabolites, including methoxylated flavonoids (*m/z* ∼365.11), flavonoids/isoflavones (*m/z* ∼381.09) and flavonoid glycosides (*m/z* ∼543.13), and a diacylglycerol (DG) lipid species (*m/z* ∼617.49) as the major compounds underlying the observed localisation patterns.

These spatial patterns demonstrate clear chemical differentiation within the grain, reflecting tissue-specific molecular organisation that is not resolved by bulk omics approaches. Furthermore, these distributions indicate functional partitioning of metabolite classes, with lipid species associated with embryo/aleurone tissues and flavonoid-related metabolites enriched in the crease region. The co-localisation of flavonoid metabolites (*m/z* ∼365.11 and ∼381.09) to the crease region indicates a spatially confined flavonoid-related metabolic domain, while the broader distribution of flavonoid glycosides (*m/z* ∼543.14) in the endosperm is consistent with downstream storage or compartmentalisation of flavonoid-related metabolites.

## Discussion

### Heat impacts are developmentally dependent on timing

Previous studies showed that sustained heat events during grain filling reduce grain size by shortening the grain-filling period, thereby limiting assimilate accumulation during development (Barrero et al. 2020). By contrast, this study reveals profound impacts of 48-h heatwaves prior to grain filling, producing a molecular memory that remains evident in the grain at the mid-filling stage. Heat exposure around anthesis impairs grain set rather than grain filling. Paradoxically, reduced grain number decreases sink competition among those florets that survive and provides them with increased assimilate supply, as proposed by the competition hierarchy among grain siblings (Shen et al. 2023). Thus, heat stress during the developmental stages around anthesis affects grain establishment through mechanisms entirely distinct from those operating during grain filling.

Illustrating the impact of subtle differences in the timing of heatwaves, exposure to heat at the trinucleate stage caused a large reduction in grain set and substantial changes in mature-grain amino acid profiles, whereas heat at the binucleate stage (pre-anthesis) and immediately post-anthesis stages (3–6 d after anthesis) had comparatively smaller effects on grain traits, despite inducing measurable metabolic changes. For instance, proline significantly accumulated in mature grain but only after exposure to heat at TN, consistent with its role as an osmoprotectant and antioxidant involved in cellular responses to heat stress (Tiwari 2024). Proline accumulation, widescale decline in free amino acids and a distinct metabolomic profile observed at the TN stage, compared with other developmental stages, indicate that the metabolic response to heat stress was highly dependent on the timing of exposure. This pattern contrasts with *higher* free amino acid content in developing wheat grains subjected to heat stress during grain filling (Wang et al. 2018), highlighting the marked interaction between heat events and developmental timing on metabolic responses. These findings demonstrate that sensitivity to temperatures above 36°C persists beyond meiosis (Masoomi-Aladizgeh et al. 2020) and varies qualitatively throughout the developmental stages that lead to grain maturation. The molecular signatures and grain traits induced by heat stress shortly before or at anthesis are a previously under-appreciated window of reproductive heat sensitivity in wheat.

Meiosis is widely recognised as one of the most heat-sensitive stages of pollen development, primarily due to disruption of recombination and meiotic restitution processes (Endo et al. 2009; Draeger and Moore 2017; Begcy et al. 2019; Masoomi-Aladizgeh et al. 2020; Masoomi-Aladizgeh et al. 2024; De Storme and Geelen 2020; De Jaeger-Braet et al. 2022). However, our data indicate that reproductive heat sensitivity is not confined to meiosis. While heat stress during the binucleate stage did not reduce pollen viability at anthesis, heat stress imposed only two days later at the trinucleate stage resulted in abortion of approximately half the florets. This sudden onset of heat sensitivity highlights the vulnerability of reproductive processes occurring during anthesis, including pollination and fertilisation events (Liu et al. 2023). Similar findings were reported in maize in which high temperatures did not reduce pollen viability. Nonetheless, inhibition of sperm cell development and transport into pollen tubes reduced fertilisation rate and grain set (Li et al. 2024).

Post-anthesis heat exposure (3 DAA than at 6 DAA) did not significantly affect grain number or weight in this study but induced notable changes in amino acid composition. Because grain biomass was not affected by post-anthesis heat exposure, the observed amino acid changes suggest altered metabolism during early grain development rather than impaired grain growth, consistent with evidence that heat can radically disrupt metabolism without necessarily reducing final grain mass (Girousse 2023). These findings demonstrate that heat exposure at pre-anthesis and anthesis affects both grain set and composition. In contrast, heat exposure during the early post-anthesis stages examined here (3–6 DAA) primarily altered grain biochemical properties without significantly affecting grain yield. This contrasts with reports that heat stress during later grain-filling stages reduces starch content and grain weight (Zhao et al. 2022), highlighting the strong influence of developmental timing on heat-stress outcomes.

### Decoupling of molecular reprogramming and phenotypic outcomes

A key finding of this study is the decoupling of molecular responses to heat stress and traits of mature grain. While pre-anthesis heat (BN) induced broader epigenetic and transcriptional changes, including more extensive DNA methylation variation and broader transcriptional reprogramming, heat during anthesis (TN) resulted in stronger downstream impacts on grain set and composition. Therefore, the magnitude of molecular reprogramming resulting from a 48-h heatwave did not predict grain phenotype. Hence, we conclude that developmental stages prior to grain filling determine how stress-induced molecular changes are translated into grain properties. The contrasting responses observed at BN and TN indicate that the magnitude of molecular reprogramming alone is insufficient to predict phenotypic outcomes; rather, the developmental context in which those responses occur is equally important. Similar uncoupling between biological processes and agronomic outcomes has been reported for complex cereal traits, including grain number and grain size (Yin et al. 2026).

Different omics approaches often show weak or variable correlations owing to the hierarchical and context-dependent nature of biological regulation (Masoomi-Aladizgeh et al. 2021; Ryan and Robards 2006). This complexity extends to whole-plant phenotypes which can only be predicted through the layered application of omics technologies (do Amaral and Souza 2017). Consistent with this, the strong molecular responses at the methylation and transcriptome levels at IMG-BNH did not translate into a notable phenotypic effect in grain. Despite TN having 82% fewer heat-responsive transcripts than BN, substantial floret abortion and changes in grain composition were evident in mature grain following heat exposure at TN but not at BN. These findings also align with systems-level perspectives, suggesting that higher-order organisational states, such as developmental stage, influence how molecular responses are translated into phenotypic outcomes (Noble 2012).

Overall, these findings reveal that developmental stage is a key determinant of how stress-induced molecular responses are translated into phenotypic outcomes, highlighting the importance for breeders in considering developmental stages as they select for heat tolerance in novel wheat cultivars.

### Tissue-specific organisation of heat responses

Spatial RNA-seq revealed extensive transcriptional heterogeneity within wheat grains in the mid-filling stage, identifying seven distinct functional domains in tissues defined by classical anatomy. Transcript clusters have been further resolved *within* functional groups in grain filling under optimal conditions (Millsteed et al. 2025; Li et al. 2025) but, to date, there is no information on how heat stress during reproductive stages, a major environmental stressor in wheat, affects these domains. This study confirms extensive, spatially organised heat-induced transcriptional reprogramming, with some specific grain regions exhibiting particularly strong transcriptional responses, while others remained comparatively stable. Notably, the modified aleurone, nucellar projection and vascular-associated tissues exhibited some of the strongest transcriptional responses to heat stress, with pronounced changes in defence, proteostasis and regulatory pathways. In contrast, embryo-associated regions showed a qualitatively different response, characterised by the maintenance of translational and biosynthetic signatures. These patterns indicate that heat-induced transcriptional reprogramming was concentrated within specific grain domains rather than being uniformly distributed across tissues. Similar spatially heterogeneous responses have been observed in developing wheat grains exposed to post-anthesis heat stress, where embryo-associated and nutrient-transfer tissues exhibited distinct transcriptional responses to elevated temperature (Wang et al. 2024). Our results extend these observations by demonstrating that brief heat events imposed at closely spaced reproductive stages (BN and TN) generate distinct spatial transcriptional programs. Unlike previous studies that primarily examined heat responses during grain filling, our analyses demonstrated that the spatial organisation of heat responses changes over a narrow developmental window spanning pre-anthesis and anthesis. Pre-anthesis and anthesis heat stress generated distinct tissue-specific molecular programs despite their close developmental proximity, highlighting the exceptional sensitivity of developing grain to the timing of heat exposure. These findings underline the interaction between spatial organisation and developmental timing in shaping responses to brief heatwaves and show that tissues within developing grain have distinctive responses that reflect their respective functional roles.

The spatial organisation revealed by transcriptomics was further supported by DESI-MSI, which revealed distinct distributions of flavonoid- and lipid-related metabolites. Flavonoid-associated ions were enriched in the crease and modified aleurone regions, whereas lipid-associated species accumulated predominantly within embryo and aleurone tissues. These findings are consistent with previous evidence that wheat grains exhibit strong spatial functional organisation, whereby distinct anatomical regions contribute different protective, metabolic and storage-related functions (Jerkovic et al. 2010).

Together, transcriptomic and mass spectrometry imaging data reveal that molecular responses to heatwaves imposed shortly before or at anthesis are expressed in a highly localised manner, with distinct tissues exhibiting specialised transcriptional and metabolic responses that collectively shape grain adaptation to elevated temperature.

### A conceptual model underlying increased individual grain weight

Although the precise mechanisms responsible for the increased individual grain weight observed following anthesis heat stress remain unresolved, our results support a model in which reduced grain number decreases sink competition among surviving grains, thereby increasing resource availability during subsequent grain development (Fig. 12). Although assimilate partitioning was not measured directly, this interpretation is consistent with source–sink models in wheat in which reductions in sink number increase resource availability for surviving grains (Shen et al. 2023). However, hormonal regulation might also contribute, as floret abortion may trigger cell division and enhance assimilate unloading in the remaining viable sinks.

**Figure 12.**
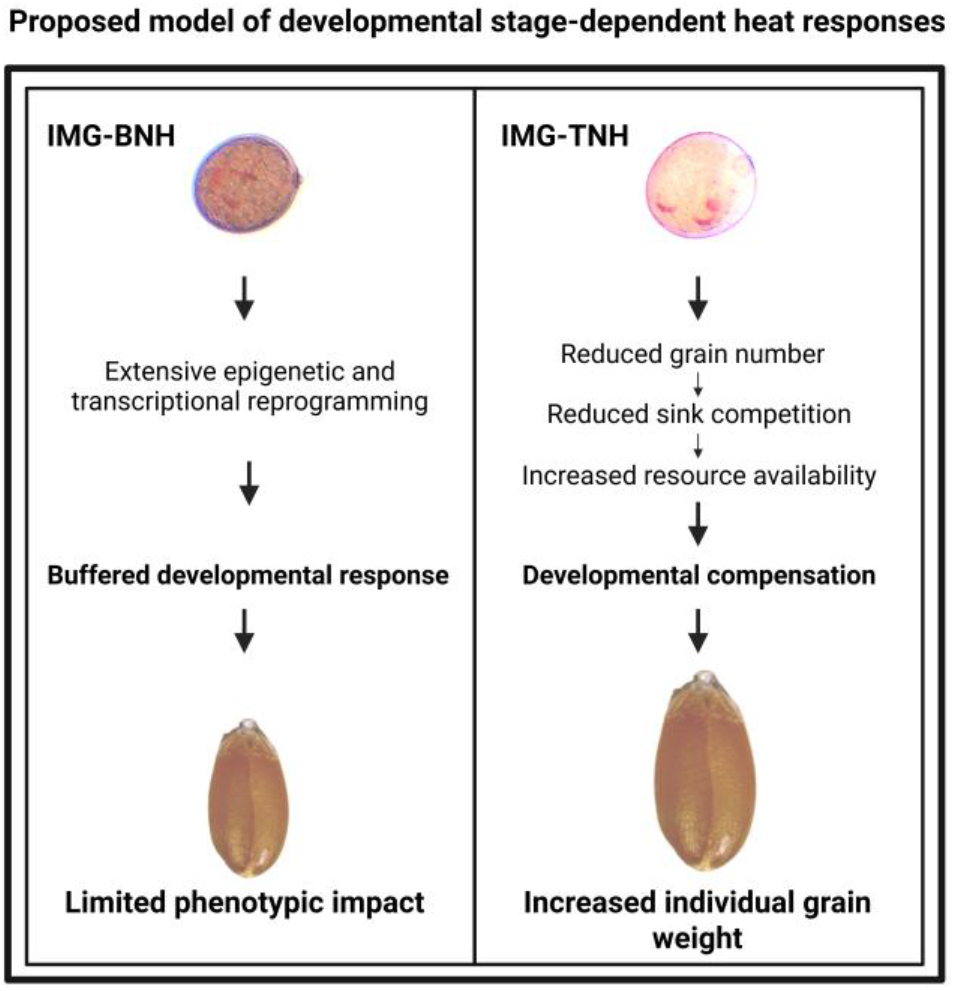
Proposed model of developmental-stage-dependent heat stress responses in wheat grain development. Heat stress (36/29°C for 48 h) at the binucleate stage (BNH) induces extensive molecular reprogramming but results in a buffered developmental response and limited phenotypic impact. In contrast, the elevated temperature at anthesis (TNH) reduces grain number, decreases sink competition and increases resource availability, supporting sustained biosynthetic activity, developmental compensation and increased individual grain weight.

Spatial transcriptomic analyses provide a potential molecular rationale for this response. Embryo-associated tissues, including the scutellum and plumule, retained signatures of active translation and energy metabolism, including elevated expression of ribosomal proteins (*RPS4, RPL21, RPL36B* and *RPL29A*), the translation elongation factor *EEF1D* and ATP synthase-associated genes (*ATPC, ATPd* and *ATPO*). These pathways are fundamental to cellular growth and biosynthetic activity, suggesting that embryo-associated tissues retained developmental capacity despite heat stress, as nucleotide metabolism underpins RNA synthesis, protein synthesis and cellular energy transfer during plant development (Zrenner et al. 2006). In contrast, the modified aleurone, nucellar projection and vascular-associated tissues exhibited more extensive stress-associated reprogramming, including suppression of genes involved in defence (*PR4B, THI13* and *DEF1*), proteostasis (*AMPD, ACX1* and *POX2*), transcriptional regulation (*MADS58, MADS8* and *MADS29*), carbohydrate metabolism (*AGAL, E13D* and *TPS*) and transport (*HA1* and *VATC*). These patterns suggest a spatial partitioning of responses in which embryo-associated domains retain signatures of active growth and development, while transfer-associated tissues undergo broader stress adaptation.

Metabolomic analyses revealed further broad metabolic reprogramming, including alterations in primary carbon metabolism, nucleotide metabolites and stress-associated compounds, indicating shifts in metabolic priorities during grain development. DESI-MSI further demonstrated strong spatial compartmentalisation of flavonoid- and lipid-associated metabolites across grain tissues. Together with the observed epigenetic reprogramming, these findings support a model in which heat-induced reductions in grain number are accompanied by coordinated molecular compensation within surviving grains. The preservation of translational and energetic activity in embryo-associated tissues (He et al. 2015) may contribute to sustained assimilate utilisation and biomass accumulation, thereby supporting increased individual grain weight. Similar increases in grain weight have been associated with altered developmental regulatory programs and enhanced cell proliferation in cereals (Yan et al. 2024), supporting the possibility that coordinated developmental compensation contributes to the enhanced growth of surviving grains following anthesis heat stress.

## Conclusion

This study demonstrates that the timing of heat exposure during reproductive development critically determines molecular reprogramming and its downstream effects on grain yield and quality. Importantly, we show that the magnitude of molecular responses does *not* necessarily predict grain traits, revealing a developmental decoupling of molecular reprogramming with grain yield and properties. Instead, developmental stage determines whether molecular changes play a role in acclimation against heat or propagate damage throughout the grain-filling stage. The identification of anthesis as a key window of post-meiotic sensitivity highlights an important target for improving heat resilience in wheat. More broadly, the integration of spatial transcriptomics with multi-omics profiling provides a powerful framework for understanding how environmental stress reshapes developmental trajectories and ultimately influences crop productivity under climate change.

## Acknowledgment

We gratefully acknowledge financial support for this research from the Bruce Veness Chandler Fund, including a fellowship for F.M.-A. We acknowledge the Spatial Pan-Omics Initiative, at the Charles Perkins Centre, University of Sydney, for access to facilities, expertise and infrastructure supporting the spatial transcriptomic and spatial metabolomics components of this work. The authors are grateful to Arjina Shrestha for technical assistance with plant growth experiments conducted in the Controlled Environment Facilities (CEF). We also thank all staff and collaborators who contributed to sample preparation, data acquisition and analytical support.

## Author contributions

F.M.-A. conceived the study, designed and performed experiments, conducted data analysis and integration, interpreted results, and wrote the manuscript. T.H.A assisted with spatial RNA-seq experiments. L.-E.Q. assisted with metabolomics experiments and analysis. Y.A. implemented computational pipelines and performed data processing. C.M. assisted with micro-CT experiments and 3D reconstruction. S.L., A.I.B., E.J, S.D. and M.W. contributed to sample preparation and data acquisition. K.B., B.C., A.K., R.T., D.K.Y.T., and B.J.A provided conceptual input and critical feedback. T.H.R conceived and supervised the study, contributed to data interpretation, and secured funding. All co-authors contributed to the revision of the manuscript.

## Supplementary information

Supplementary Figures S1–S6 and Supplementary Data S1–S2 are available with this article.

## Data availability statement

The datasets generated and analysed during the current study will be deposited in public repositories and made available upon publication. Accession numbers will be provided in the final published article.

## Supplementary figures

**Figure S1.**
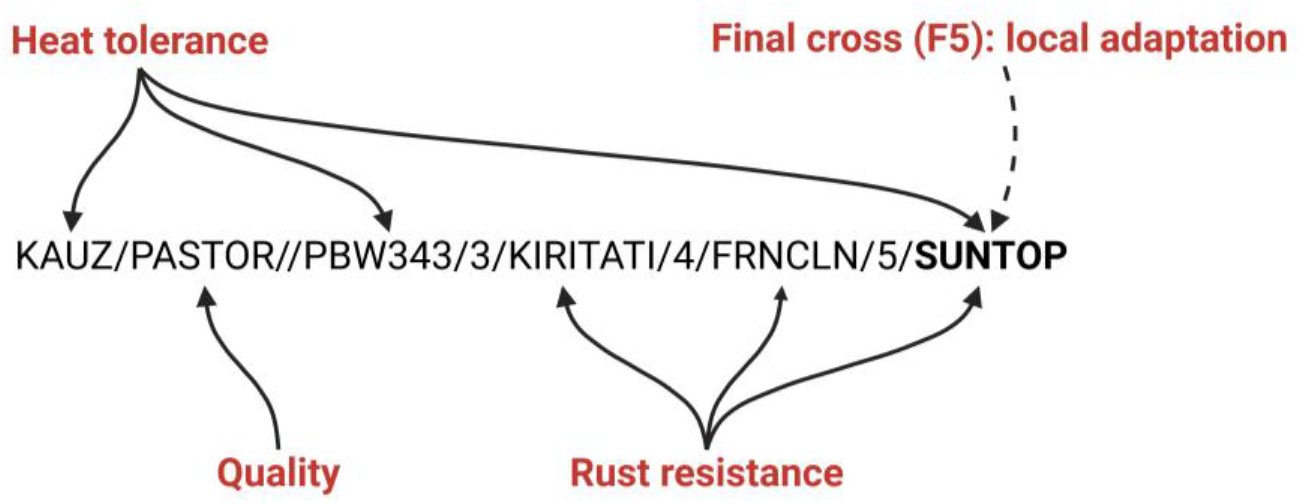
Pedigree of PBI13505 showing contributions from the CIMMYT-derived line KAUZ/PASTOR//PBW343/3/KIRITATI/4/FRNCLN, selected in Mexico for plant height, yield performance under heat stress, and disease resistance. This parental line combines heat tolerance from Kauz and PBW343, grain-quality attributes from Pastor, and durable minor-gene rust resistance from Kiritati and Francolin. The final cross (F5) was made with the locally adapted Australian cultivar Suntop, which contributed 50% of the pedigree and provides triple rust resistance and intermediate heat tolerance. The aim of the cross was to combine international sources of heat tolerance with local adaptation while further diversifying the rust-resistance background.

**Figure S2.**
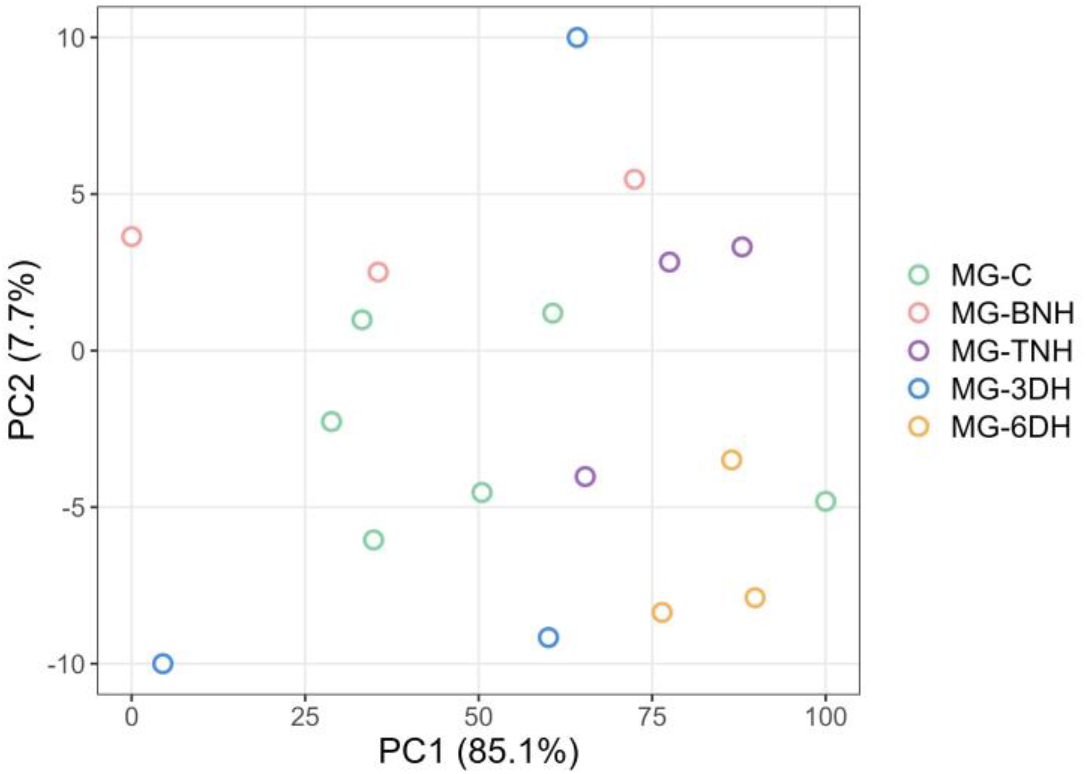
PCA of NIR spectra from mature grains under developmental heat treatments, showing scores for PC1 (85.1% of variance) versus PC2 (7.7% of variance). Treatments show partial separation, with some clustering of MG-TNH and MG-6DH samples, and greater dispersion in MG-C, MG-BNH and MG-3DH.

**Figure S3.**
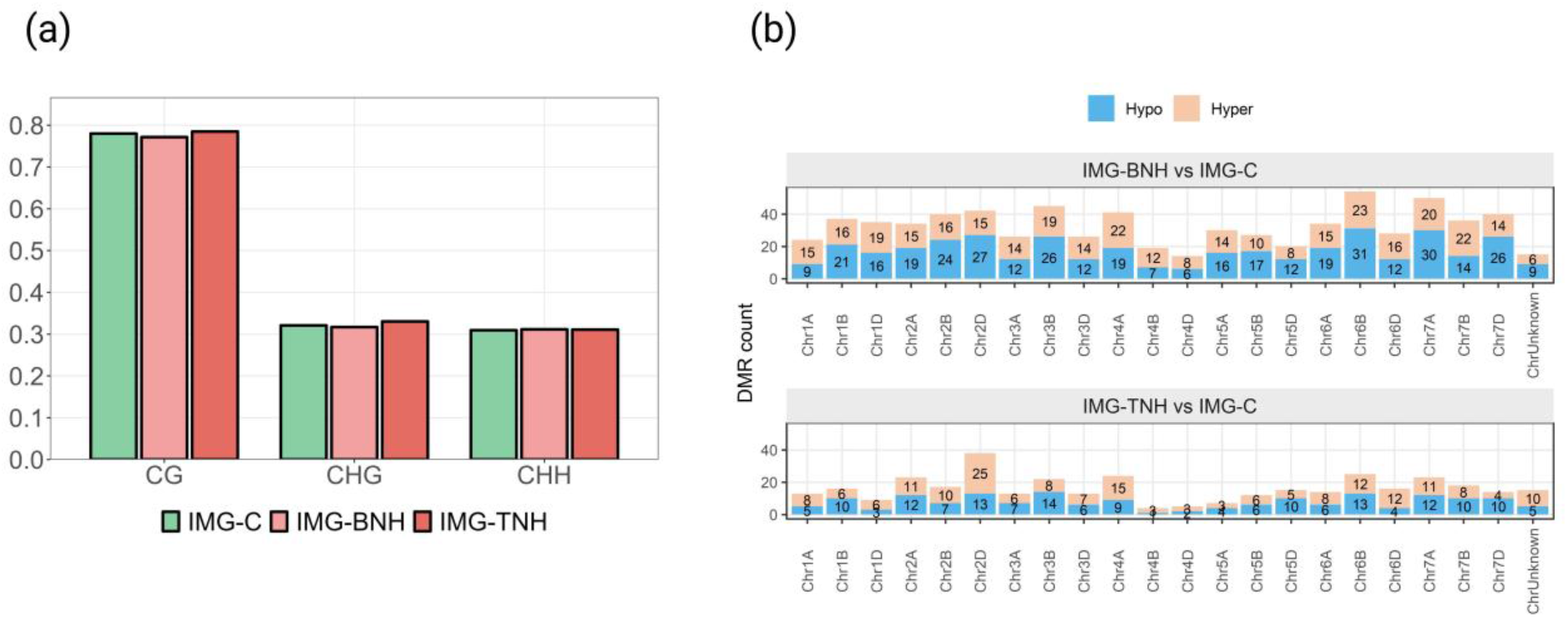
Effects of short-term heat stress (36/29 ℃ for 48 h) at defined developmental stages on methylation patterns. (a) The similar global methylation values across IMG-C, IMG-BNH, and IMG-TNH indicate that heat treatments did not induce broad epigenomic reprogramming. (b) Chromosomal distribution of DMRs (methylation difference ≥ 15%, q ≤ 0.01) is uneven across the A, B, and D subgenomes, with distinct stage-specific hotspots.

**Figure S4.**
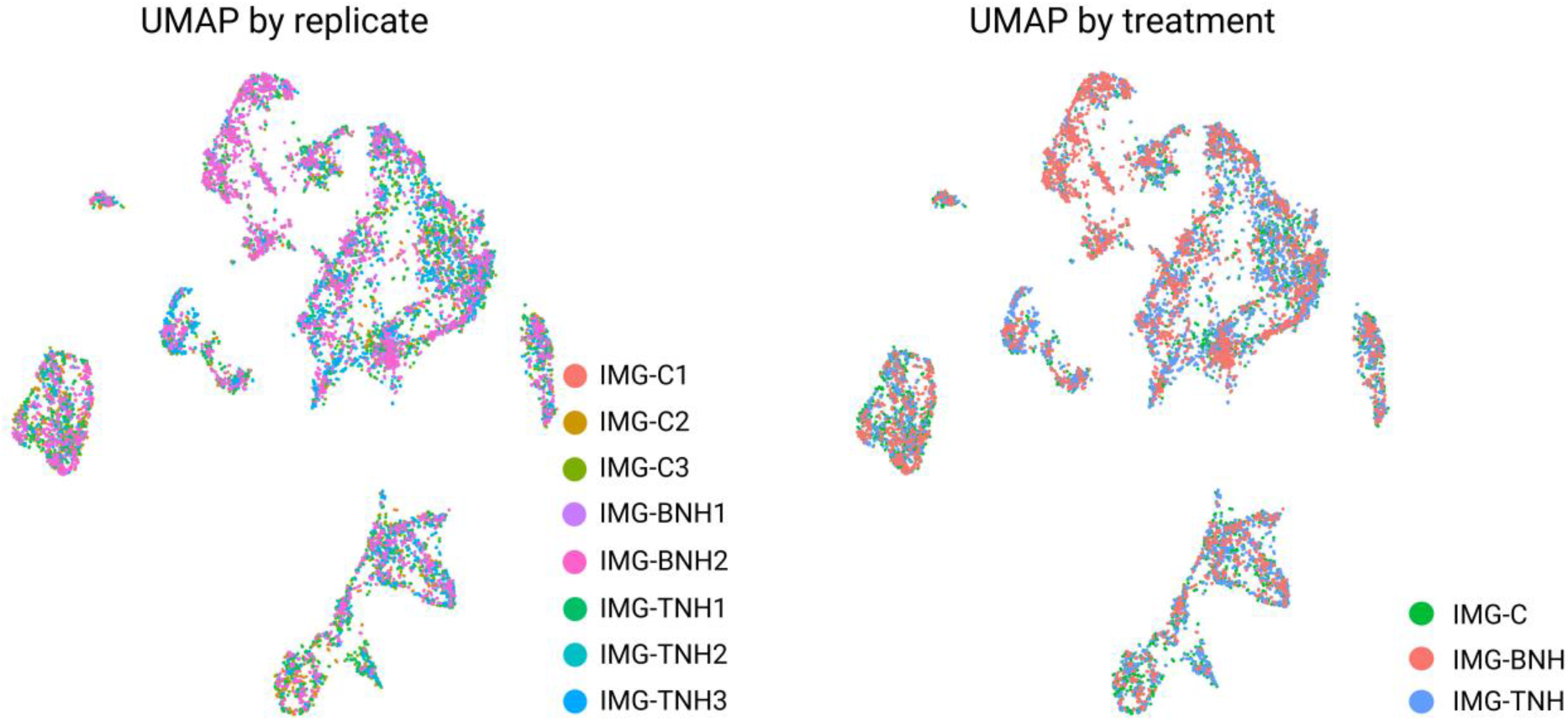
UMAP visualisation of integrated spatial transcriptomic profiles coloured by replicate and treatment, demonstrating effective integration across samples and indicating that overall transcriptional structure is independent of replicate and treatment effects.

**Figure S5.**
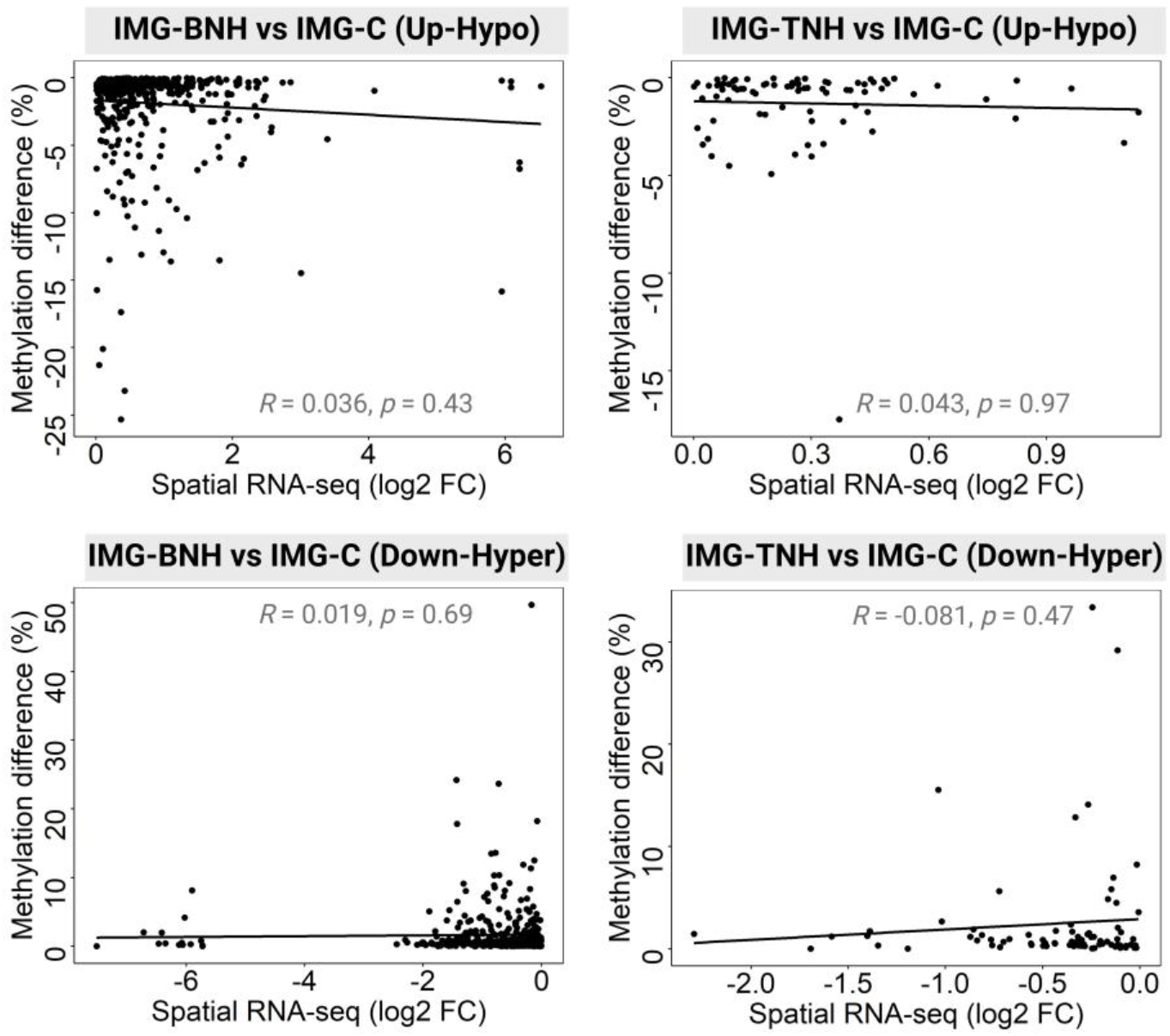
Effects of short-term heat stress (36/29 ℃ for 48 h) at defined developmental stages on the relationship between transcriptomic responses and DNA methylation. Scatter plots showing the relationship between methylation difference (%) and spatial RNA-seq (log2 fold change) for IMG-BNH vs IMG-C and IMG-TNH vs IMG-C, arranged by direction of change (up-regulated–hypo-methylated and down-regulated–hyper-methylated). No thresholds were applied for fold change or methylation difference.

**Figure S6.**
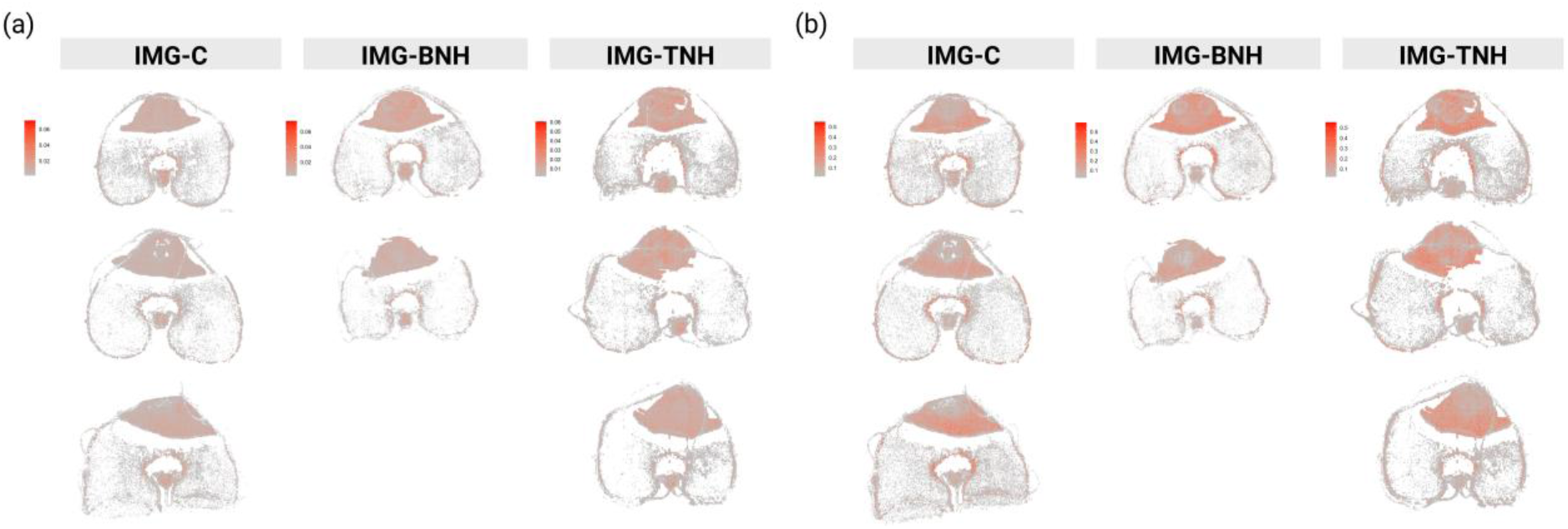
Spatial distribution of global and marker-based stress-response programs across all biological replicates. (a) Spatial distribution of the global-derived stress-response program across all biological replicates from control (IMG-C), IMG-BNH and IMG-TNH samples. (b) Spatial distribution of the marker-derived stress-response program across the same sections.

